# Design, Amino-Acid-Derived Analogues of FTY-720 and OSU-2S as PP2A Activators: Structure–Activity Relationships and Identification of Lead Compound XML-3D

**DOI:** 10.64898/2026.10.12.758679

**Authors:** Rui Yang, Noura Srour, Pankajavalli Thirugnanasambandam, Shabber Mohammed, Megan E. Roderick, Santhanam Ramasamy, Ann Ventura, Enming Xing, Jeffrey Cheng, Xiaolin Cheng, Aiden Tamulonis, Dharani Mayilsamy, Anuvrat Sarkar, Lalit Sehgal, Natarajan Muthusamy, Pui Kai Li

## Abstract

Protein phosphatase 2A (PP2A) is a critical regulator of oncogenic signaling pathways, and pharmacological reactivation of PP2A through disruption of inhibitory protein interactions has emerged as a promising anticancer strategy. Although the sphingosine analogue FTY-720 and its derivative OSU-2S can restore PP2A activity, further structural diversification is needed to improve potency and expand structure–activity understanding. Here, we report the design, synthesis, and biological evaluation of three series of amino-acid-derived analogues incorporating tyrosine, homotyrosine, and tryptophan scaffolds. Docking models were used qualitatively to guide exploration of head-group functionality, hydrophobic tail variation, and stereochemical configuration; these models were hypothesis-generating and were not experimentally validated. Modular synthetic routes enabled efficient preparation of structurally diverse compounds from naturally occurring starting materials. Biological evaluation in leukemia and lymphoma cell lines revealed that hydrophobic tail length strongly influenced cytotoxic activity, while tryptophan-derived analogues were generally less potent than tyrosine-based scaffolds. Shortening the hydrophobic tail did not improve activity, contrary to the initial docking-based hypothesis. Stereochemical inversion at the amino alcohol head group yielded a lead compound, XML-3D, which demonstrated approximately two-fold enhanced cytotoxicity relative to FTY-720 and OSU-2S across leukemia cell lines (IC₅₀ = 4.1, 1.8, and 1.5 μM in MV4-11, HL-60, and MEC-1 cells, respectively) and showed broader activity in mantle cell lymphoma lines (IC₅₀ = 530 nM, 970 nM, and 110 nM in Jeko-1, Mino-1, and UPN1 cells). XML-3D induced increased PP2A phosphatase activity in cellular assays. Co-immunoprecipitation and pharmacological inhibition experiments were consistent with modulation of the PP2A/SET regulatory axis; however, direct binding measurements were not performed, and the mechanistic interpretation remains provisional. These findings suggest preliminary structure–activity relationships for amino-acid-derived PP2A modulators and identify stereochemical optimization as a productive direction for future development. Ongoing work aims to define molecular target engagement and refine mechanistic understanding to guide next-generation design.

## 1. Introduction

Protein phosphorylation, mediated by kinases and phosphatases, is a vital cellular regulatory mechanism involved in fundamental processes including cell cycle control, growth, differentiation, metabolism, and apoptosis. (1,2) Perturbations in phosphorylation signaling are implicated in numerous diseases, including cancer, diabetes, and neurological disorders. (3,4) Through phosphorylation, protein kinases relay signals from extracellular stimuli to intracellular responses by modulating protein interactions, localization, and conformational states, ultimately influencing gene transcription and protein turnover. (1) Accordingly, kinase inhibition has emerged as a successful therapeutic strategy, and more than 80 kinase inhibitors have received FDA approval for cancer treatment. (5) In contrast, signal termination and pathway modulation are governed by protein phosphatases, which are classified into four major classes based on catalytic mechanism and substrate specificity: serine/threonine phosphatases (PSPs), tyrosine phosphatases (PTPs), dual-specificity phosphatases, and histidine phosphatases. (6) Among the PSPs, the phosphoprotein phosphatase (PPP) family includes seven human enzymes: PP1, PP2A, PP2B, PP4, PP5, PP6, and PP7. (7–9) Of these, PP2A is the most abundant and plays a central role in regulating oncogenic signaling.

Protein phosphatase 2A (PP2A) is a heterotrimeric enzyme composed of scaffolding (A), regulatory (B), and catalytic (C) subunits. (10) The A and C subunits form a core complex that associates with diverse B subunits to generate functionally distinct holoenzymes. The A subunit mediates assembly and substrate positioning, while the B subunit governs spatial and temporal specificity. (11) Both A and C subunits exist as α and β isoforms with high sequence homology, whereas over 26 structurally diverse B-subunit isoforms are encoded by multiple gene families. (6,10,12)

PP2A regulates multiple oncogenic signaling networks, including the PI3K/Akt and MAPK pathways. In the PI3K/Akt pathway, activation of PI3K leads to PIP3 formation and subsequent phosphorylation of Akt at Thr308 and Ser473 by PDK1 and mTORC2. (13,14) PP2A counteracts this activation through dephosphorylation of these residues, thereby suppressing proliferative signaling. (15,16) Similarly, PP2A attenuates MAPK signaling by dephosphorylating Raf kinases and related effectors. (17,18)

Dysregulation of PP2A is observed in diverse pathological states including neurodegeneration, cardiovascular disease, and cancer. (6) In oncology, PP2A functions as a tumor suppressor, yet its activity is frequently diminished through mutations, altered subunit expression, or upregulation of endogenous inhibitory proteins. (19–22) These mechanisms rarely eliminate PP2A expression entirely but instead attenuate its activity, suggesting that pharmacological reactivation represents a viable therapeutic strategy.

Efforts to target PP2A have therefore focused on indirect activation approaches, particularly disruption of inhibitory protein interactions. Two prominent endogenous inhibitors—SET (I2PP2A) and CIP2A—directly bind and suppress PP2A activity. (23,24) Elevated SET expression has been reported in multiple hematologic malignancies, including acute myeloid leukemia (AML) (25) and chronic lymphocytic leukemia (CLL), (26) as well as in solid tumors such as non-small cell lung cancer, (27) colorectal cancer, (20,28) and hepatocellular carcinoma. (29) Genetic or pharmacological interference with SET restores PP2A signaling and suppresses oncogenic phosphorylation cascades. (30,31)

Ceramide and related sphingolipids have been shown to disrupt SET-PP2A interactions. (32–34) One clinically established derivative is FTY720 (fingolimod), an immunomodulatory agent approved for multiple sclerosis treatment (Figure 1). Its immunosuppressive activity arises from phosphorylation and agonism of sphingosine-1-phosphate receptors, leading to lymphocyte sequestration. (35)

**Figure 1.**
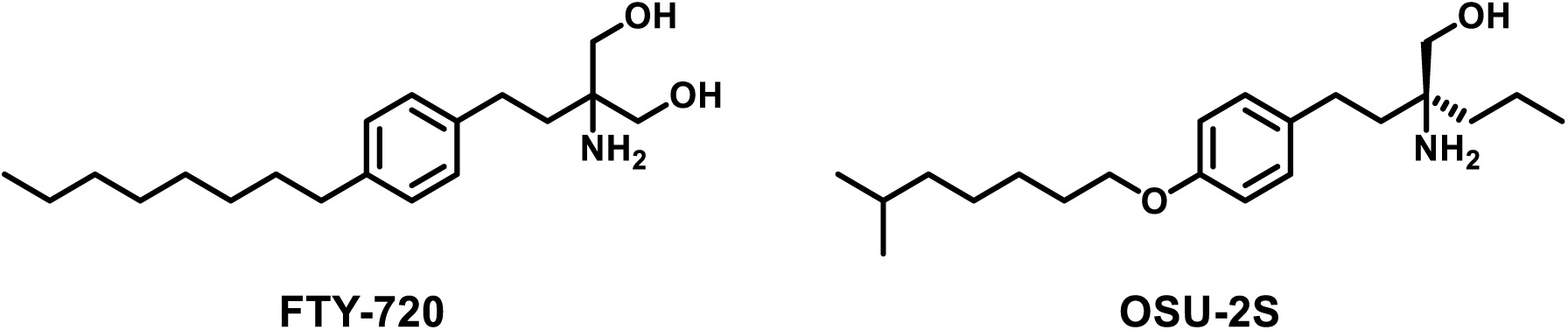
Structures of FTY-720 and OSU-2S

**Figure 2.**
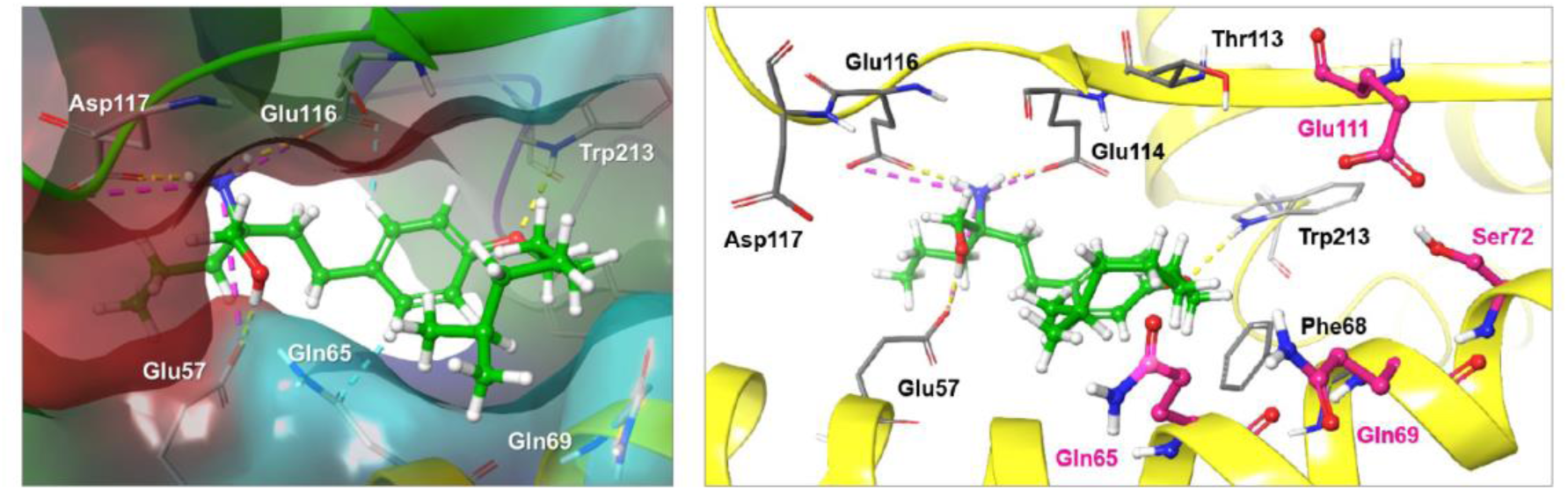
Docking pose of OSU-2S to SET

Importantly, FTY720 also demonstrates antitumor activity in several cancer models independent of receptor phosphorylation. (36–40) These effects have been associated with disruption of SET-mediated inhibition of PP2A and downstream suppression of PI3K/Akt and MAPK signaling. (38,39,41) Subsequent medicinal chemistry efforts sought to retain PP2A-modulating activity while reducing receptor-mediated immunosuppression. Structure-activity investigations of aminoethanol analogues established positions amenable to modification within the scaffold. (42) This led to development of OSU-2S, a non-immunosuppressive analogue capable of activating PP2A and inducing cytotoxicity in hematologic malignancies. (43) Further scaffold diversification demonstrated that stereochemical and lipophilic modifications influence PP2A-modulating activity and cellular responses. (44,45) These studies collectively illustrate the feasibility of tuning pharmacological properties within this chemical class while preserving functional signaling effects.

Despite these advances, systematic exploration of amino-acid-derived scaffolds that leverage synthetic accessibility and stereochemical diversity remains limited. In particular, it is unclear how the amino acid core (tyrosine vs. homotyrosine vs. tryptophan), hydrophobic tail length, and stereochemistry at the amino alcohol head group jointly influence PP2A activation and cytotoxicity. Accordingly, the present study investigates tyrosine-, homotyrosine-, and tryptophan-based derivatives designed to expand chemical diversity while maintaining key pharmacophoric features associated with PP2A activation. We report the design, synthesis, and biological evaluation of three compound series, identify hydrophobic tail length as a major determinant of potency, and demonstrate that stereochemical inversion at the head group yields a lead compound (XML-3D) with improved cytotoxicity and PP2A activation relative to FTY-720 and OSU-2S.

## 2. Design and Synthesis of FTY-720 and OSU-2S Derivatives as PP2A Activators

### 2.1 Design of FTY-720 and OSU-2S Derivatives

Saddoughi et al. identified a potential binding region for FTY-720 on SET through molecular modeling and mutagenesis, implicating residues K209 and Y122 in ligand interaction. (40) To gain insight into possible interactions of OSU-2S with SET, OSU-2S was docked into the same region of the SET crystal structure (PDB 2E50).

Docking simulations generated a putative binding pose that was used solely to guide hypothesis generation rather than to infer definitive binding interactions. These models were not experimentally validated, and the docking results should be interpreted qualitatively.

Three observations emerged from this model:

1. The hydrophobic tail adopted a bent conformation extending away from polar residues (Glu111, Ser72, Gln69, Gln65).
2. The propyl substituent of the head group showed minimal predicted interaction.
3. The amino and hydroxyl groups were positioned to form hydrogen bonds with Glu116/Glu114 and Glu57, respectively.

These observations were interpreted qualitatively to guide structural modification; however, docking models alone cannot establish binding mode or affinity, and no direct binding measurements were performed in this study.

Based on these insights, three modification strategies were explored (Figure 3):

**1. Tail modification.** We explored whether shortening the tail length and incorporating polar functionality could improve interaction with proximal polar residues.
**2. Head group simplification.** Removal of the propyl group while retaining amino and hydroxyl functionalities was pursued to streamline synthesis and preserve hydrogen-bonding capacity.
**3. Core scaffold diversification.** Because OSU-2S derives from L-homotyrosine, alternative amino-acid cores (L-tyrosine and L-tryptophan) were evaluated to expand structural diversity and probe steric/electronic tolerance.

**Figure 3.**
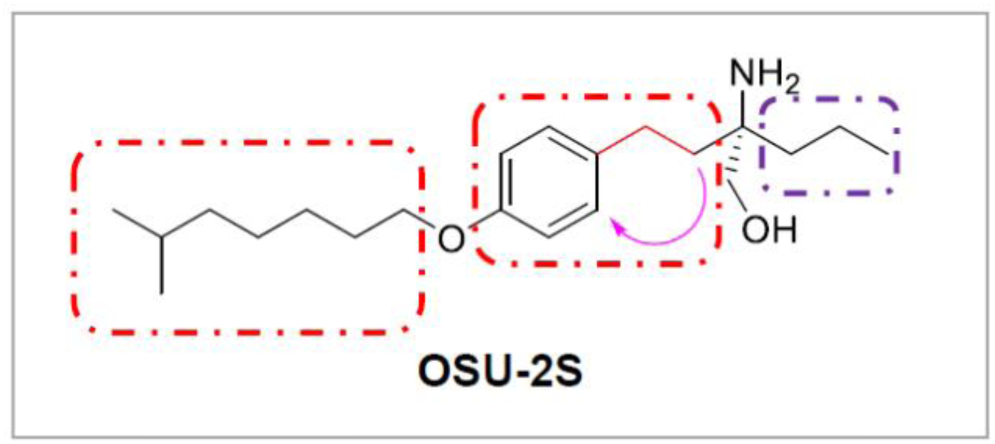
Modification strategies for OSU-2S

Collectively, these strategies were intended to explore structure–activity relationships rather than to validate a specific docking-derived binding hypothesis. As described below, the experimental data did not support the initial hypothesis that tail shortening would enhance activity; instead, analogues retaining extended hydrophobic substituents exhibited superior cellular potency.

We synthesized three compound series (Figure 4):

- L-tyrosine derivatives (XML-1–3, 10–14)
- L-homotyrosine derivatives (XML-4–6)
- L-tryptophan derivatives (XML-7–9)

**Figure 4.**
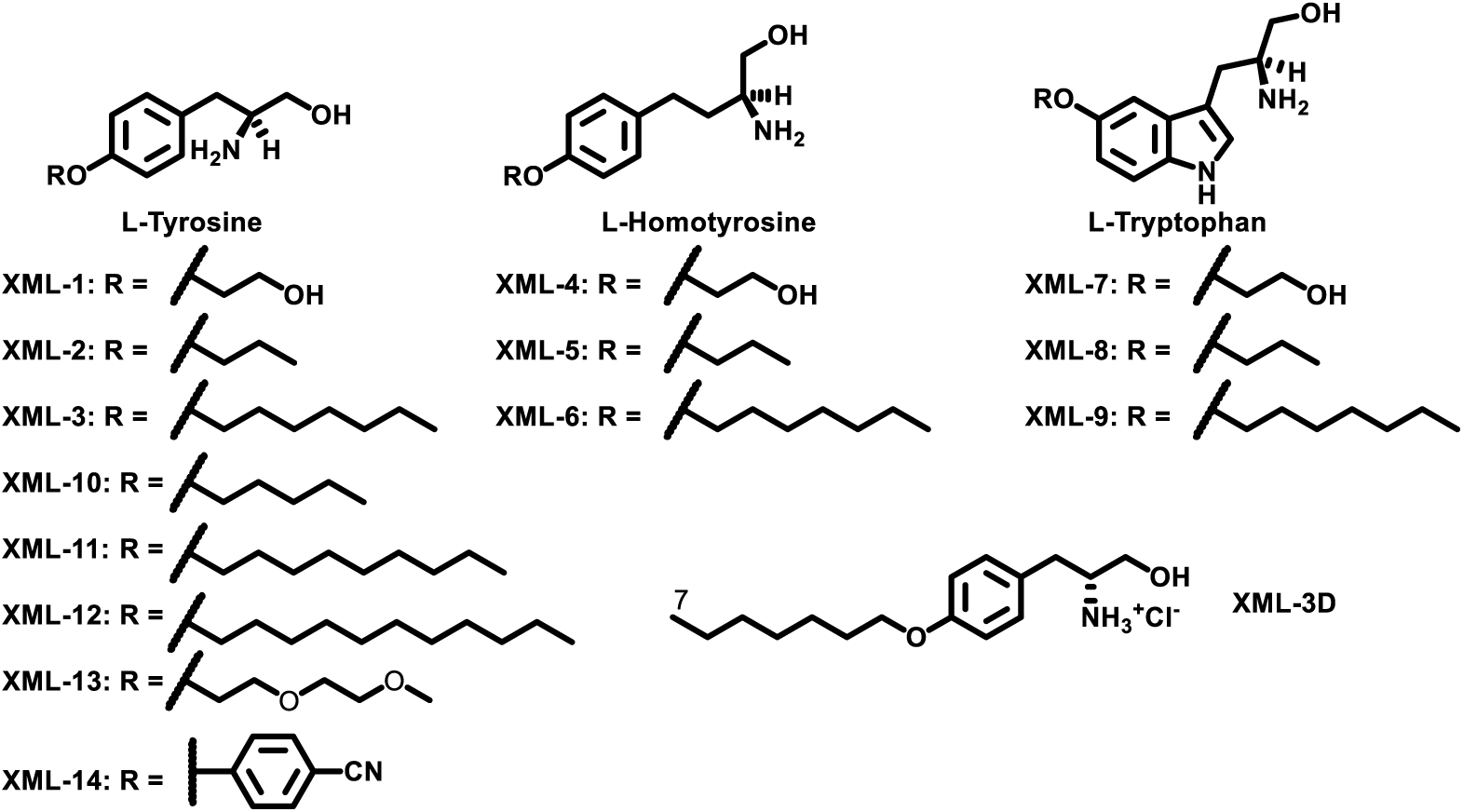
Structures of OSU-2S derivatives

XML-1–9 incorporated three tail types (hydroxyethyl, propyl, heptyl), enabling comparison of polarity and length effects. XML-10–14 varied hydrophobic tail size more broadly to assess steric contributions. A D-enantiomer (XML-3D) was included to evaluate stereochemical effects on activity.

This library design allowed systematic exploration of tail length, core scaffold identity, and stereochemistry within a synthetically accessible framework. Compared with FTY-720, OSU-2S, and AAL(S), whose syntheses require enantiomeric resolution or chiral intermediates, our approach utilized naturally occurring amino acids as starting materials. This enabled convergent preparation through shared intermediates and avoided post-synthetic resolution steps.

### 2.2 Synthesis

The L-tyrosine series was prepared beginning from L-tyrosine methyl ester (Scheme 1). Boc protection of the amino group was followed by alkylation or aryl substitution of the phenolic oxygen. Reduction of the methyl ester with NaBH₄/LiCl afforded the corresponding alcohol, and acidic Boc removal yielded the final ammonium salts.

Reaction sequences proceeded reproducibly across substrates, enabling preparation of multiple analogues using a unified protocol.

**Scheme 1.**
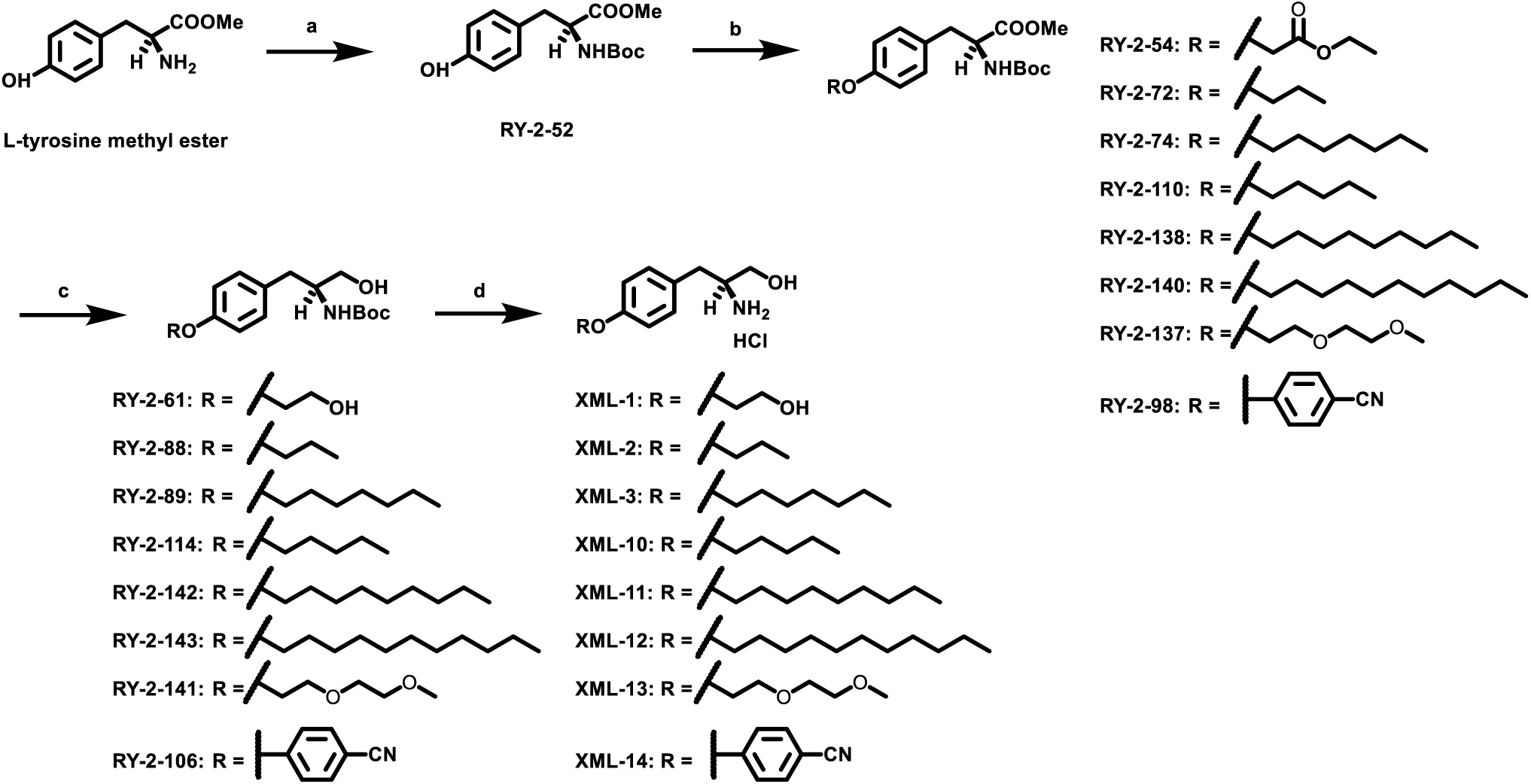
Syntheses of XML 1–3 and 10–14. (a) di-tert-butyl dicarbonate, sodium bicarbonate, MeOH, 79%; (b) alkyl bromide/4-fluorobenzonitrile, potassium carbonate, DMF, 80 °C, overnight, 23–96%; (c) sodium borohydride, lithium chloride, EtOH/THF (1:1), 60 °C, overnight, 41–93%; (d) 4N HCl in 1,4-dioxane, 2 h, 58– 88%.

L-homotyrosine and L-tryptophan derivatives were synthesized analogously (Schemes 2 and 3) via esterification, protection, tail installation, ester reduction, and deprotection.

**Scheme 2.**
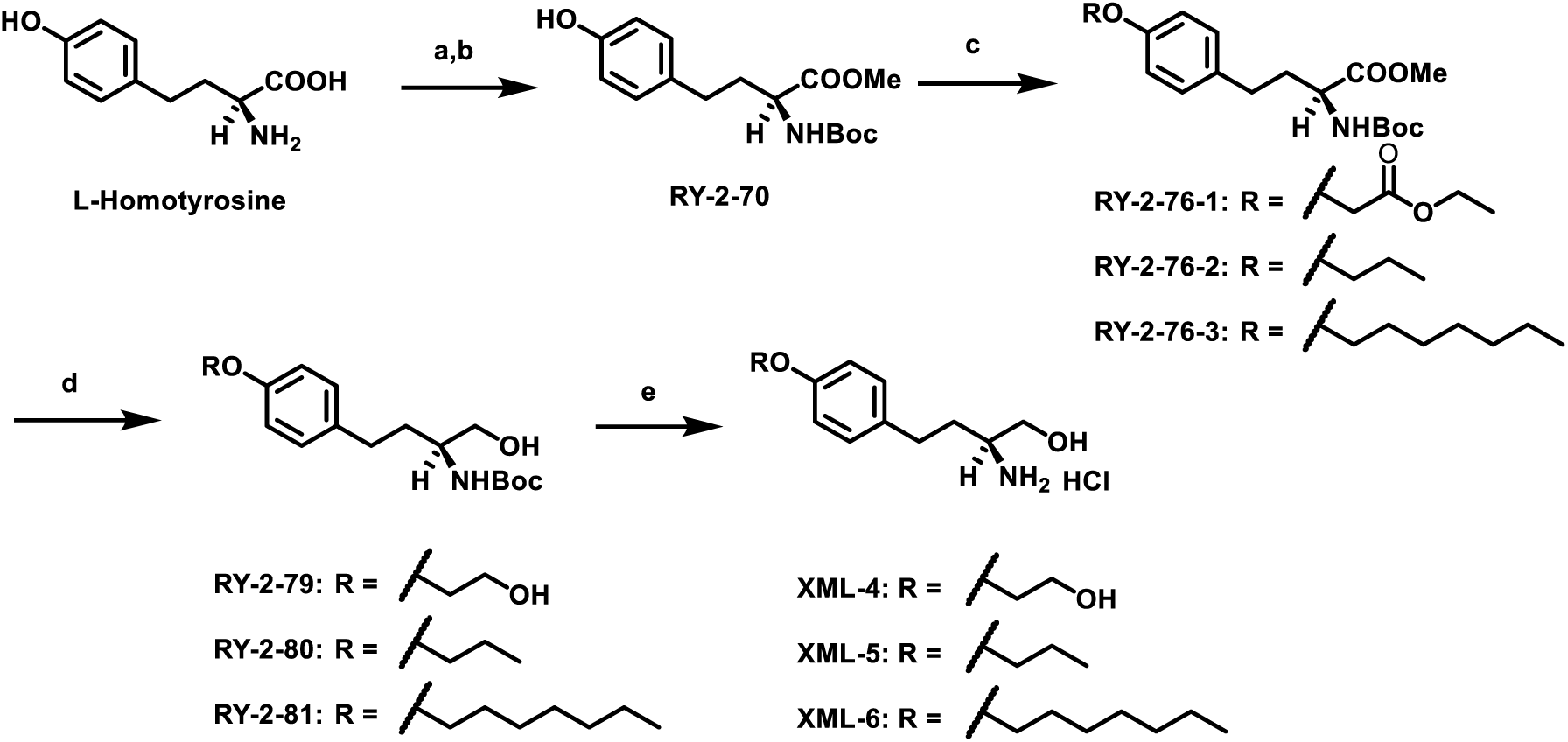
XML 4–6. Reagents and conditions: (a) thionyl chloride, MeOH, 60 °C; (b) di-tert-butyl dicarbonate, sodium bicarbonate, MeOH, 56% over two steps; (c) alkyl bromide, potassium carbonate, DMF, 80 °C, overnight, 88–98%; (d) sodium borohydride, lithium chloride, EtOH/THF (1:1), 60 °C, overnight, 38–75%; (e) 4N HCl in 1,4-dioxane, 2 h, 85–97%.

**Scheme 3.**
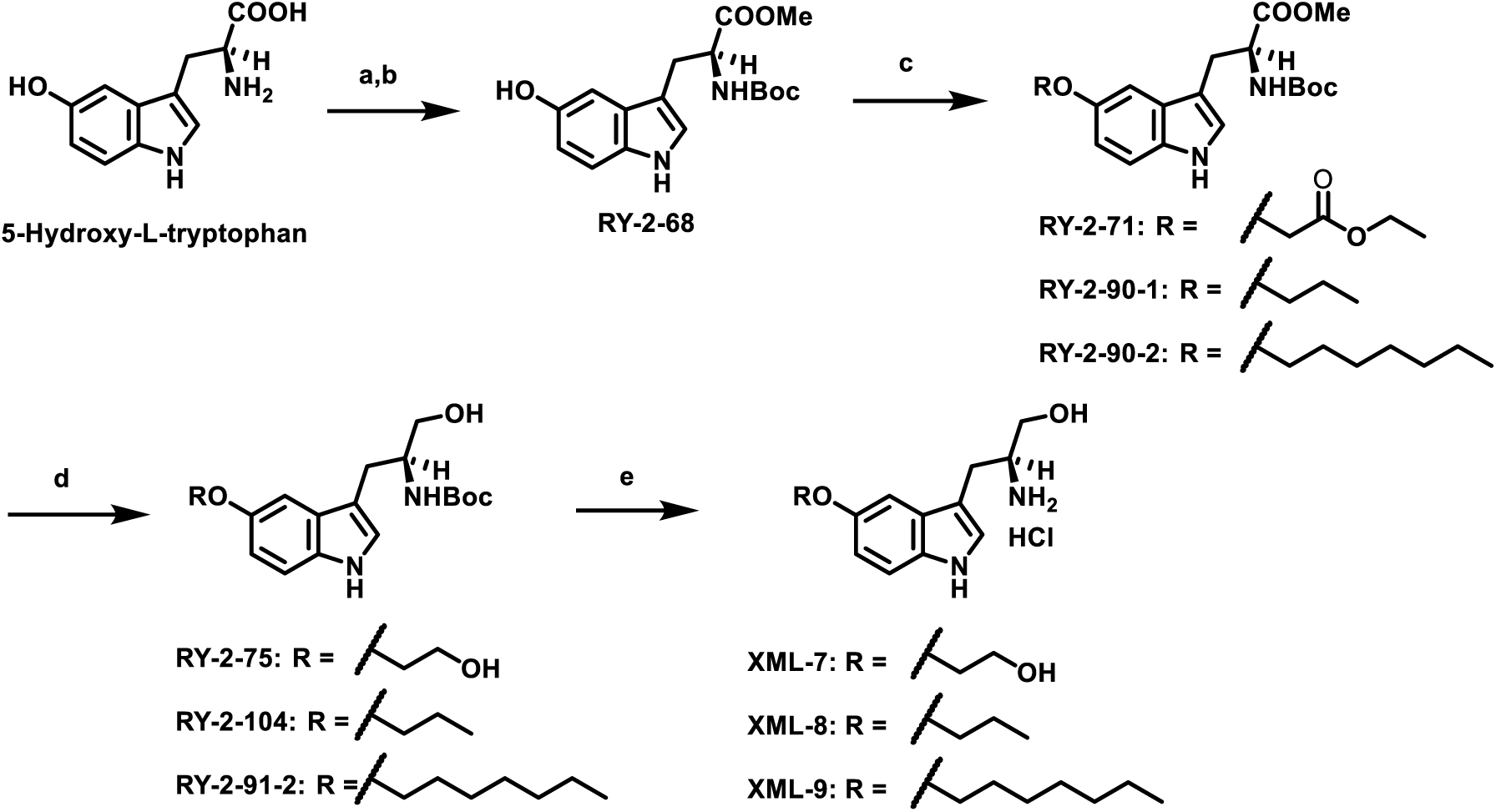
Synthesis of XML 7–9. Reagents and conditions: (a) thionyl chloride, MeOH, 60 °C; (b) di-tert-butyl dicarbonate, sodium bicarbonate, MeOH, 45% over two steps; (c) alkyl bromide, potassium carbonate, DMF, 80 °C, overnight, 34–74%; (d) sodium borohydride, lithium chloride, EtOH/THF (1:1), 60 °C, overnight, 48–96%; (e) 4N HCl in 1,4-dioxane, 2 h, 58–88%.

XML-3D was generated using D-tyrosine as starting material (Scheme 4).

**Scheme 4.**
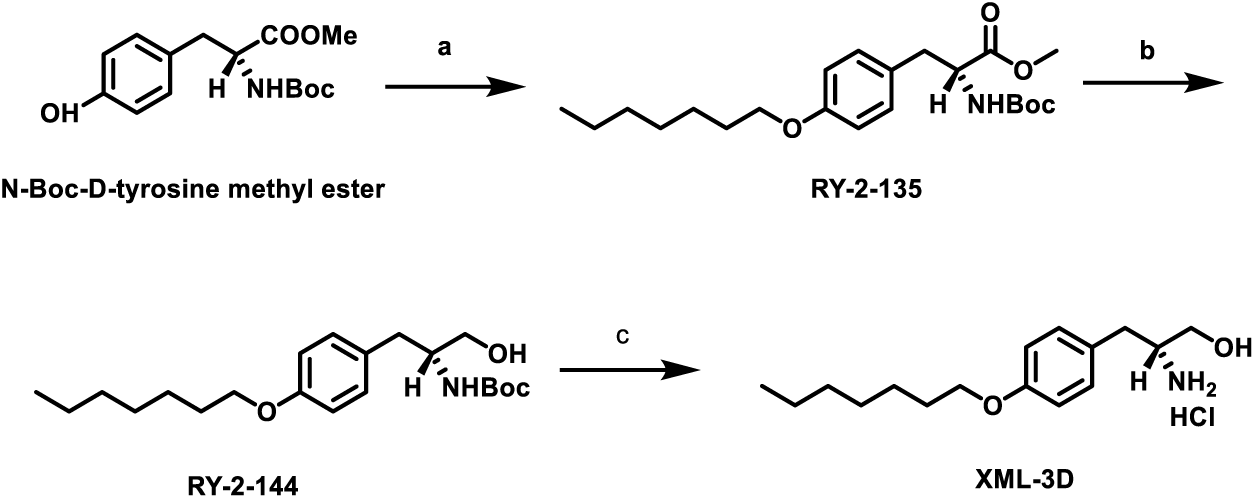
Synthesis of XML-3D. Reagents and conditions: (a) 1-bromoheptane, potassium carbonate, DMF, 80 °C, overnight, 89%; (b) sodium borohydride, lithium chloride, EtOH/THF (1:1), 60 °C, overnight, 93%; (c) 4N HCl in 1,4-dioxane, 2 h, 70%.

All final compounds were characterized by ¹H NMR, ¹³C NMR, and HRMS-ESI, and purity was assessed by UPLC using two independent gradient methods (A1/B1 and A2/B2). All compounds used for biological testing exhibited ≥85% purity by UPLC; the lowest purity observed was 86.54% for XML-10 (A2/B2 method). Purity data for all final compounds are provided in the Supplemental Information. Given that all compounds tested were ≥85% pure, impurity contributions to biological activity are considered unlikely, although this possibility cannot be entirely excluded for XML-10.

Overall, the modular nature of the synthetic strategy enabled efficient generation of structurally diverse analogues suitable for biological evaluation.

### Biological Results

The cytotoxicity of XML compounds was evaluated in human acute myeloid leukemia (AML; MV4-11, HL-60), chronic lymphocytic leukemia (CLL; MEC-1), and mantle cell lymphoma (MCL; Jeko-1 and Mino) cell lines using an MTS viability assay. Cytotoxicity data are summarized in Table 1, with only compounds exhibiting measurable IC₅₀ values reported. Compounds with IC₅₀ > 50 μM were considered inactive in this assay. All compounds tested were ≥85% pure by UPLC (see Supplemental Information), and impurity contributions to activity are therefore considered unlikely, although this possibility cannot be entirely excluded for XML-10 (86.54% purity).

**Table 1.** Cytotoxicity of XML series compounds.

| Compound | MV4-11 $IC_{50}$ (μM) | HL-60 $IC_{50}$ (μM) | MEC-1 $IC_{50}$ (μM) |
| --- | --- | --- | --- |
| XML-1 | >50 | >50 | >50 |
| XML-2 | >50 | >50 | >50 |
| XML-3 | 7.02 | 6.26 | 7.06 |
| XML-4 | >50 | >50 | >50 |
| XML-5 | >50 | >50 | >50 |
| XML-6 | 5.01 | 4.11 | 7.73 |
| XML-7 | >50 | >50 | >50 |
| XML-8 | >50 | >50 | >50 |
| XML-9 | 9.23 | 9.62 | 15.43 |
| XML-10 | 29.62 | 27.20 | >30 |
| XML-11 | 8.40 | 7.10 | 5.50 |
| XML-12 | 7.80 | 6.70 | 4.30 |

| Compound | MV4-11 IC<sub>50</sub> (μM) | HL-60 IC<sub>50</sub> (μM) | MEC-1 IC<sub>50</sub> (μM) |
| --- | --- | --- | --- |
| XML-14 | 40.30 | 40.78 | 81.30 |
| XML-3D | 4.10 | 1.80 | 1.50 |
| FTY-720 | 10.30 | 7.70 | 6.90 |
| OSU-2S | 6.70 | 6.40 | 5.70 |

Note: IC<sub>50</sub> values were determined from non-linear regression curves using GraphPad Prism 9. All experiments were performed with three technical replicates. Independent biological replicates were not performed, and standard deviations are therefore not reported. Compounds with IC<sub>50</sub> > 50 μM were considered inactive.

Compounds XML-3 and XML-6, which possess tail lengths comparable to OSU-2S, demonstrated IC₅₀ values similar to those of FTY-720 and OSU-2S. Specifically, XML-3 exhibited IC₅₀ values of 7.02, 6.26, and 7.06 μM in MV4-11, HL-60, and MEC-1 cells, respectively, compared with 10.3, 7.7, and 6.9 μM for FTY-720 and 6.7, 6.4, and 5.7 μM for OSU-2S. These data indicate that substitution of the L-homotyrosine core with L-tyrosine does not substantially alter cytotoxic activity within this scaffold class. In contrast, XML-9 displayed reduced potency (IC₅₀ = 9.23, 9.62, and 15.43 μM in MV4-11, HL-60, and MEC-1 cells, respectively), suggesting that incorporation of an L-tryptophan core may be less favorable for activity. The reduced potency of tryptophan-derived analogues may reflect steric bulk or altered electronic properties of the indole ring; however, this hypothesis was not experimentally tested.

Derivatives bearing shorter alkyl tails (XML-1, XML-2, XML-4, XML-5, XML-7, XML-8, and XML-10) showed little to no cytotoxic effect (IC₅₀ > 50 μM in most cell lines), whereas analogues with extended hydrophobic chains (XML-11 and XML-12) retained activity (IC₅₀ = 5.5–8.4 μM across cell lines). Collectively, these observations indicate that hydrophobic tail length is a major determinant of cellular potency in this series.

Notably, experimental trends did not support the initial hypothesis that tail shortening would enhance activity; instead, analogues retaining extended hydrophobic substituents exhibited superior cellular potency.

The activity of XML-11 and XML-12 (IC₅₀ = 4.3–8.4 μM) was comparable to that of OSU-2S (IC₅₀ = 5.7–6.7 μM), suggesting that these compounds do not offer a substantial potency advantage over the reference compound. However, they may offer altered physicochemical properties or selectivity profiles that warrant further investigation.

Notably, stereochemical inversion at the head group significantly affected potency. XML-3D, the D-enantiomer of XML-3, demonstrated approximately two-fold greater cytotoxic activity than FTY-720 and OSU-2S across leukemia cell lines (IC₅₀ = 4.1, 1.8, and 1.5 μM in MV4-11, HL-60, and MEC-1 cells, respectively, compared with 6.7–10.3 μM for FTY-720 and 5.7–6.7 μM for OSU-2S). This finding suggests that stereochemical orientation of the amino alcohol moiety influences biological engagement and highlights chirality as an important parameter for optimization. However, the mechanism underlying this stereochemical effect (e.g., improved binding affinity, metabolic stability, or cellular uptake) was not investigated in this study.

Consistent with viability data, PP2A activity assays performed in MV4-11 cells showed that XML-3D produced stronger phosphatase activation than OSU-2S (approximately 7-fold vs. 4-fold increase at 5 μM and 2.5 μM, respectively; Figure 5A). These results supported prioritization of XML-3D for mechanistic characterization. However, statistical significance was not assessed due to the limited number of replicates (n = 3 technical replicates), and these data should be interpreted as indicative rather than definitive.

**Figure 5.**
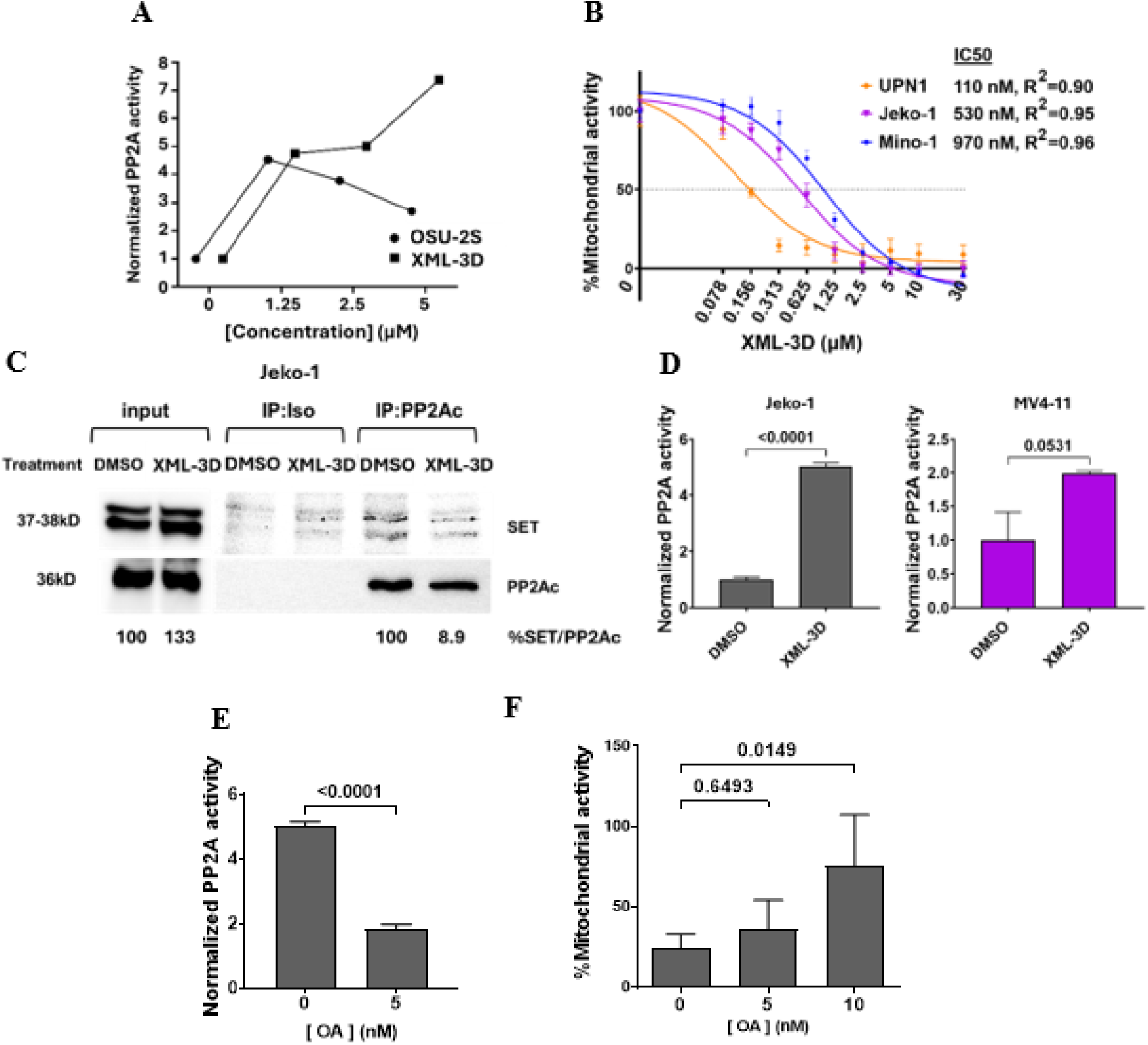
OSU-2S derivative XML-3D activates PP2A in leukemia and lymphoma cell lines. A) PP2A activity assay on MV4-11 cell line after treatment with different doses of XML-3D, OSU-2S, or untreated for 4 h. B) MTS cytotoxicity assay in 3 independent mantle cell lymphoma cell lines (Jeko-1, Mino-1, UPN1) treated with different doses of XML-3D or DMSO for 24 h. C) Jeko-1 cells were treated with 1 μM XML-3D or DMSO for 6 h followed by immunoprecipitation with anti-PP2Ac or isotype control antibody before probing for SET and PP2Ac by Western blot. D) PP2A activity assay in Jeko-1 cell line and MV4-11 cell line after treatment with 5 μM XML-3D or DMSO for 6 h and 4 h, respectively. E) PP2A activity assay on Jeko-1 cell line after treatment with 5 μM XML-3D or DMSO for 6 h in the presence or absence of pretreatment with 5 nM or 10 nM OA (PP2Ac inhibitor) for 2 h, appropriately normalized to DMSO or DMSO with OA. F) MTS viability assay on Jeko-1 cell line after treatment with 1 μM of XML-3D or DMSO for 12 h in the presence or absence of pretreatment with 5 nM or 10 nM PP2A inhibitor okadaic acid (OA) for 2 h, appropriately normalized to DMSO, DMSO with 5 nM OA, or DMSO with 10 nM OA. Note that statistics are representative of 3 technical replicates (A, B, D, E) or 4 technical replicates (F) for each cell line. Independent biological replicates were not performed for co-immunoprecipitation (C).

Given reports that FTY-720 derivatives exhibit activity in lymphoma models, XML-3D was evaluated in mantle cell lymphoma lines. The compound induced dose-dependent cytotoxicity in Jeko-1 (IC₅₀ = 530 nM), Mino-1 (IC₅₀ = 970 nM), and UPN1 (IC₅₀ = 110 nM) cells, demonstrating broader activity beyond leukemia systems (Figure 5B). These IC₅₀ values are substantially lower than those observed in leukemia cell lines, suggesting that XML-3D may exhibit cell-line-specific activity. However, the mechanism underlying this differential sensitivity was not investigated.

To examine the potential mechanism, co-immunoprecipitation studies assessed interaction between the PP2A catalytic subunit (PP2Ac) and its endogenous inhibitor SET. XML-3D treatment reduced the SET/PP2Ac ratio relative to vehicle controls in Jeko-1 cells, consistent with decreased complex association (Figure 5C). The SET/PP2Ac ratio was quantified as 100% for DMSO-treated cells and 8.9% for XML-3D-treated cells in the IP:PP2Ac lane. However, these results are from a single experiment (n = 1), and independent biological replicates were not performed. Statistical significance was therefore not assessed, and these data should be interpreted as preliminary. In line with co-immunoprecipitation experiments, XML-3D increased PP2A phosphatase activity in Jeko-1 and MV4-11 cell lines (Figure 5D). The increase in PP2A activity was approximately 4.7-fold in Jeko-1 cells (p < 0.0001) and 2.0-fold in MV4-11 cells (p = 0.0531) relative to DMSO controls. While the Jeko-1 result reached statistical significance, the MV4-11 result was borderline, likely due to the limited number of replicates (n = 3 technical replicates).

Functional relevance of PP2A activation was evaluated using the phosphatase inhibitor okadaic acid (OA). OA pretreatment largely reversed XML-3D-induced phosphatase activation and partially rescued cytotoxicity in a dose-dependent manner (Figure 5E, F). Specifically, XML-3D-induced PP2A activity was reduced from approximately 5-fold above baseline to approximately 2-fold above baseline in the presence of 5 nM OA (p < 0.0001; Figure 5E). Similarly, XML-3D-induced cytotoxicity was partially rescued by OA pretreatment (Figure 5F), with cell viability increasing from approximately 30% to approximately 45% in the presence of 5 nM OA and to approximately 80% in the presence of 10 nM OA. However, the rescue was incomplete, suggesting that PP2A-independent mechanisms may also contribute to XML-3D-induced cytotoxicity. These findings support a PP2A-dependent component underlying the biological effects of XML-3D, but they do not exclude additional mechanisms.

It should be noted that okadaic acid inhibits both PP1 and PP2A, with selectivity for PP2A at low nanomolar concentrations. The concentrations used in this study (5–10 nM) are consistent with preferential PP2A inhibition; however, selectivity was not experimentally confirmed in the cell lines used. Therefore, the possibility that PP1 inhibition contributes to the observed rescue cannot be entirely excluded.

Taken together, the biological data suggest preliminary structure–activity relationships for this compound series and identify stereochemically optimized derivative XML-3D as the most active analogue evaluated. The data are consistent with a model in which XML-3D disrupts the PP2A/SET interaction, leading to increased PP2A activity and PP2A-dependent cytotoxicity. However, direct binding measurements were not performed, and the co-immunoprecipitation data are from a single experiment. Therefore, the mechanistic interpretation remains provisional and will require confirmation through future biophysical and functional characterization, including direct binding assays, SET knockdown or overexpression studies, and PP2A-specific inhibition experiments.

### Conclusions and Outlook

Protein phosphatase 2A (PP2A) plays a central role in regulating cellular proliferation and survival through suppression of multiple oncogenic signaling pathways. Functional inhibition of PP2A by endogenous proteins such as SET is frequently observed in hematologic malignancies, making pharmacological disruption of the PP2A/SET interaction an attractive therapeutic strategy. While the sphingosine analogue FTY-720 has demonstrated the ability to restore PP2A activity, its clinical utility in oncology is limited by immunosuppressive effects mediated through sphingosine-1-phosphate receptor signaling. Development of non-immunosuppressive derivatives such as OSU-2S has partially addressed this limitation, although improvements in potency remain desirable.

In the present study, three series of amino-acid-derived analogues were designed and synthesized to explore structural diversification around the OSU-2S scaffold while maintaining synthetic accessibility. The use of naturally occurring amino acids as starting materials enabled convergent preparation through shared intermediates, avoiding post-synthetic resolution steps and facilitating efficient generation of structurally diverse analogues. Docking-guided hypotheses initially suggested that shortening and polarizing the hydrophobic tail might improve interaction with SET; however, experimental cytotoxicity data did not support this prediction. Instead, analogues retaining extended hydrophobic substituents exhibited superior cellular activity, highlighting the importance of lipophilicity in maintaining functional engagement. This negative result is informative for future design: it suggests that the hydrophobic tail contributes to target engagement or cellular accumulation in a manner that is not captured by the qualitative docking model.

Structure–activity analysis indicated that tyrosine- and homotyrosine-derived cores were generally more favorable than tryptophan-based analogues. The reduced potency of tryptophan-derived compounds may reflect steric bulk or altered electronic properties of the indole ring, although this hypothesis was not experimentally tested. Similarly, the observation that XML-11 and XML-12 displayed potency comparable to OSU-2S suggests that these compounds do not offer a substantial potency advantage over the reference compound, although they may offer altered physicochemical or selectivity profiles that warrant further investigation. Furthermore, stereochemical inversion at the amino alcohol head group produced a marked increase in potency, as exemplified by compound XML-3D. This compound consistently displayed improved cytotoxic activity relative to reference molecules and induced greater PP2A activation across leukemia and lymphoma cell models. The approximately two-fold improvement in potency over FTY-720 and OSU-2S is modest but reproducible across multiple cell lines, and the substantially lower IC₅₀ values observed in mantle cell lymphoma lines (110–970 nM) suggest that XML-3D may exhibit cell-line-specific activity. The mechanism underlying this differential sensitivity was not investigated.

Mechanistic evaluation demonstrated decreased association between PP2Ac and SET and increased phosphatase activity following XML-3D treatment, effects that were attenuated by okadaic acid. These findings are consistent with a model in which XML-3D disrupts the PP2A/SET interaction, leading to increased PP2A activity and PP2A-dependent cytotoxicity. However, several limitations should be acknowledged. First, direct binding measurements were not performed, and definitive molecular engagement with SET cannot yet be established. Second, the co-immunoprecipitation data are from a single experiment (n = 1), and independent biological replicates were not performed. Third, okadaic acid inhibits both PP1 and PP2A, and selectivity for PP2A was not experimentally confirmed in the cell lines used. Fourth, statistical analyses were limited to technical replicates, and independent biological replicates were not performed for most assays. Finally, the incomplete rescue of cytotoxicity by okadaic acid suggests that PP2A-independent mechanisms may also contribute to the biological effects of XML-3D. Accordingly, the mechanistic interpretation remains provisional.

The findings of this study should be contextualized within the broader landscape of PP2A activators in development. Several small-molecule PP2A activators have been reported, including LB-100, DT-061, and iHAP1, which act through diverse mechanisms including direct PP2A activation, disruption of PP2A/inhibitor interactions, and modulation of PP2A holoenzyme assembly. (46,47) XML-3D differs from these compounds in its amino-acid-derived scaffold and its proposed mechanism of SET displacement. However, direct comparisons of potency, selectivity, and mechanism across these compound classes were not performed in this study. Future work should include head-to-head comparisons with established PP2A activators to better define the unique features of the XML series.

The drug-likeness of XML-3D was not explicitly assessed in this study. With a molecular weight of approximately 280 Da and a cLogP value predicted to be in the range of 3–4 (based on the heptyl tail and amino alcohol head group), XML-3D is expected to comply with Lipinski’s rule of five. However, experimental determination of solubility, permeability, and metabolic stability was not performed. These properties will be important to evaluate in future studies to assess the potential for in vivo evaluation.

Future work will focus on addressing the mechanistic gaps identified in this study. Specifically, biophysical approaches including surface plasmon resonance (SPR), isothermal titration calorimetry (ITC), or microscale thermophoresis (MST) will be used to measure direct binding between XML-3D and SET. Target-engagement assays, such as cellular thermal shift assays (CETSA) or drug affinity responsive target stability (DARTS), will be employed to confirm SET as the molecular target in cells. SET knockdown or overexpression studies will be performed to validate the proposed mechanism of action. PP2A-specific inhibitors or siRNA will be used to confirm that the biological effects of XML-3D are mediated through PP2A. Additionally, structural characterization of ligand–protein complexes will be pursued to refine the binding mode and guide rational design. These studies will refine structure–activity relationships and guide the development of next-generation analogues with improved potency and mechanistic definition.

Beyond mechanistic characterization, future medicinal chemistry efforts should prioritize several directions. First, the stereochemical optimization strategy that yielded XML-3D should be extended to other scaffolds within the series, including homotyrosine and tryptophan cores, to determine whether the effect of stereochemistry is generalizable. Second, the hydrophobic tail should be further diversified to explore the balance between lipophilicity and aqueous solubility, with the goal of improving drug-like properties while maintaining potency. Third, the amino alcohol head group should be modified to probe additional hydrogen-bonding interactions with SET, guided by the docking model but validated through experimental binding studies. Fourth, analogues with improved metabolic stability should be designed, particularly those resistant to oxidative metabolism of the alkyl tail. Finally, the selectivity of XML-3D for PP2A over other phosphatases, particularly PP1, should be systematically evaluated.

Collectively, the present results suggest preliminary structure–activity relationships for amino-acid-derived PP2A modulators and identify stereochemical optimization as a promising direction for continued development of PP2A-modulating agents. The lead compound XML-3D demonstrates improved potency and PP2A activation relative to FTY-720 and OSU-2S, and its activity in mantle cell lymphoma lines supports further investigation in hematologic malignancies. However, the mechanistic interpretation remains provisional due to the limitations outlined above. Addressing these limitations through future biophysical, biochemical, and pharmacological studies will be essential to establish XML-3D as a viable lead compound and to guide the rational design of next-generation PP2A activators.

## Experimental Section and Methods

### 1. General Chromatographic and Analytical Procedure

All reactions were performed at room temperature unless otherwise stated. Reagents and solvents were purchased from commercial suppliers. All air- or moisture-sensitive reactions were performed under nitrogen using anhydrous solvents and glassware. All instrumentation used for the structural characterization and confirmation was in the College of Pharmacy at The Ohio State University. ^1^H and ^13^C NMR were obtained on the Bruker Avance 400/300 MHz spectrometer Chemical shifts (*δ*) are reported in ppm and coupling constants (*J* values) are reported in Hz. High-resolution Mass Spectra-Electron Spray Ionization (HRMS-ESI) were obtained using the Thermo Q-Exactive Orbitrap with Vanquish-H UHPLC. The LC/MS analyses were obtained using the Acquity UPLC@BEH C18 column (1.7µm, 2.1 x 50 mm) on the Waters Acquity UPLC with an increasing gradient of 2−98% B (A1 = water + 0.1% formic acid, B1 = methanol + 0.1% formic acid; A2 = water + 0.1% formic acid, B2 = acetonitrile + 0.1% formic acid) over 5.0 min and then 98% B for 2.0 min and back to 2% B over 1 min at a flow rate of 0.35 mL/min. The peak detection was determined by UV absorbance at 270 nm.

RediSep gold and silver silica gel disposable flash columns, 40-60 micron and empty disposable sample load cartridges (Teledyne ISCO) were used for flash column chromatography on CombiFlash (Teledyne ISCO). Reagent grade solvents for flash column chromatography were obtained from Fisher Scientific. MilliporeSigma™ Silica Gel 60 F254 Coated Aluminum-Backed TLC Sheets were used for TLC. General compound visualization for TLC was performed under UV light (254 nm and 365 nm).

### 2.4 Synthesis of XML compounds

#### Synthesis of RY-2-52

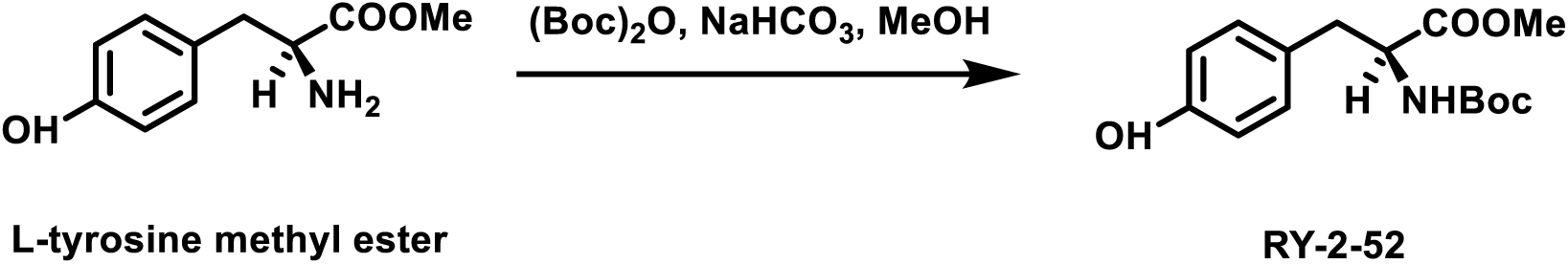

L-tyrosine methyl ester (1950 mg, 10 mmol, 1.0 eq) was dissolved in methanol (20 mL) followed by the addition of sodium bicarbonate (1680 mg, 20 mmol, 2.0 eq). A solution of Boc anhydride (2619 mg, 12 mmol, 1.2 eq) in methanol (10 mL) was added. The reaction was stirred at room temperature overnight. Upon completion, the solvent was dried under reduced pressure and the residue was re-dissolved in water (200 mL). The mixture was acidified with 1N HCl until pH = 4 and extracted with ethyl acetate (100 mL x 3). The organic phase was washed with brine (100 mL x 3) and dried over sodium sulfate. The solvent was removed under reduced pressure and the residue was subjected to flash column chromatography (25% ethyl acetate in hexane). The product was obtained as white solid with a yield of 2.3 g, 79%. ^1^H NMR (400 MHz, Chloroform-*d*) δ 7.66 (s, 1H), 6.95 (d, *J* = 8.3 Hz, 2H), 6.77 (d, *J* = 8.1 Hz, 2H), 5.23 (d, *J* = 8.4 Hz, 1H), 4.52 (dt, *J* = 8.5, 6.1 Hz, 1H), 3.68 (s, 3H), 3.07 – 2.94 (m, 1H), 1.42 (s, 9H).

#### Synthesis of RY-2-54

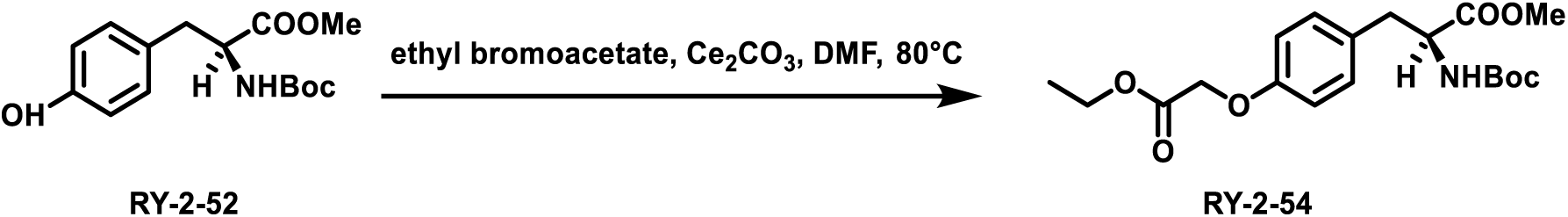

RY-2-52 (1476 mg, 5 mmol, 1.0 eq) was dissolved in DMF (20 mL) followed by the addition of ethyl bromoacetate (1002 mg, 6 mmol, 1.2 eq) and cesium carbonate (3258 mg, 10 mmol, 2.0 eq). The reaction mixture was heated at 80°C and stirred overnight. Upon completion, the reaction mixture was quenched with water (100 mL) and extracted with ethyl acetate (100 mL x 3). The organic phase was washed with brine (100 mL x 3) and dried over sodium sulfate. The solvent was removed under reduced pressure and the residue was subjected to flash column chromatography (25% ethyl acetate in hexane). The product was obtained as colorless oil with a yield of 430 mg, 23%. ^1^H NMR (400 MHz, Chloroform-*d*) δ 7.05 (d, *J* = 8.4 Hz, 2H), 6.84 (d, *J* = 8.7 Hz, 2H), 4.99 (d, *J* = 8.2 Hz, 1H), 4.54 (d, *J* = 7.7 Hz, 1H), 4.27 (q, *J* = 7.1 Hz, 2H), 3.70 (s, 3H), 3.02 (qd, *J* = 14.0, 5.9 Hz, 2H), 1.42 (s, 9H), 1.30 (t, *J* = 7.1 Hz, 3H).

#### Synthesis of RY-2-61

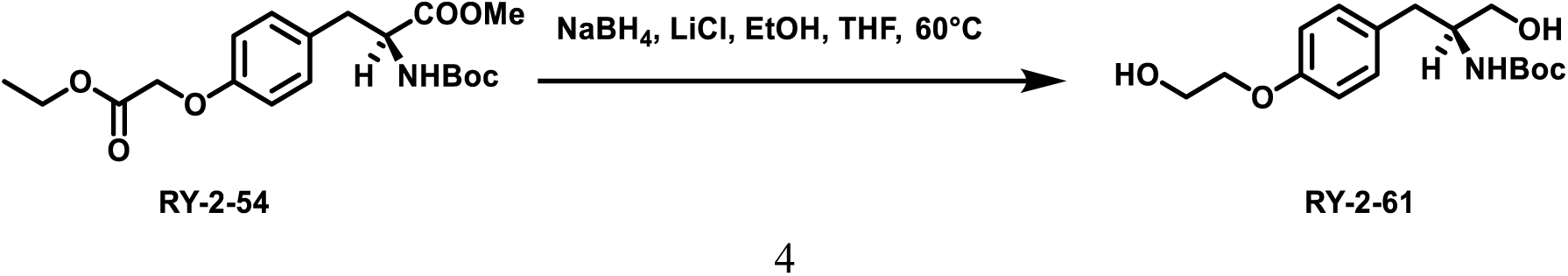

RY-2-54 (381 mg, 1 mmol, 1.0 eq) was dissolved in ethanol (10 mL) and THF (10 mL) followed by the addition of sodium borohydride (303 mg, 8 mmol, 8.0 eq) and lithium chloride (339 mg, 8 mmol, 8.0 eq). The reaction mixture was heated at 60°C and stirred overnight. Upon completion, the reaction mixture was slowly quenched with water (50 mL) and extracted with ethyl acetate (100 mL x 3). The organic phase was washed with brine (100 mL x 3) and dried over sodium sulfate. The solvent was removed under reduced pressure and the residue was subjected to flash column chromatography (90% ethyl acetate in hexane). The product was obtained as white solid with a yield of 264 mg, 85%. ^1^H NMR (300 MHz, Methanol-*d*4) δ 7.14 (d, *J* = 8.1 Hz, 2H), 6.86 (d, *J* = 8.1 Hz, 2H), 4.01 (s, 1H), 3. 87 (s, 1H), 3.73 (s, 1H), 3.51 (s, 1H), 3.02 – 2.48 (m, 3H), 1.39 (s, 9H).

#### Synthesis of XML-1

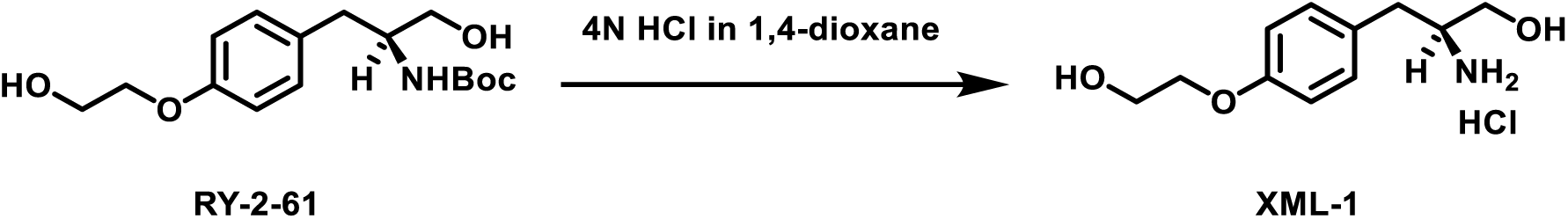

To a round-bottom flask was added RY-2-61 (200 mg, 0.6 mmol) and 4N HCl in 1,4-dioxane (5 mL). The reaction mixture was stirred for 2 h. Upon completion, the solvent was removed under reduced pressure. The residue was suspended in ethyl ether (5 mL) and filtered. The solid was washed with ethyl ether (10 mL) and dried under vacuum. The product was obtained as white solid with a yield of 100 mg, 63%. ^1^H NMR (400 MHz, Methanol-*d*4) δ 7.20 (d, *J* = 8.6 Hz, 2H), 6.94 (d, *J* = 8.7 Hz, 2H), 4.08 – 3.93 (m, 2H), 3.92 – 3.77 (m, 2H), 3.69 (dd, *J* = 11.6, 3.5 Hz, 1H), 3.52 (dd, *J* = 11.6, 6.3 Hz, 1H), 3.45 – 3.34 (m, 1H), 2.88 (d, *J* = 7.4 Hz, 2H). ^13^C NMR (101 MHz, Methanol-*d*4) δ 158.33, 129.98, 127.66, 114.65, 69.19, 60.28, 59.97, 54.56, 34.18. MS (ESI) calculated for [C11H18NO3]^+^: 212.27, found: 212.2. UPLC: tR = 2.191 min, >99% (A1/B1); tR = 1.899 min, >99% (A2/B2).

#### Synthesis of RY-2-72

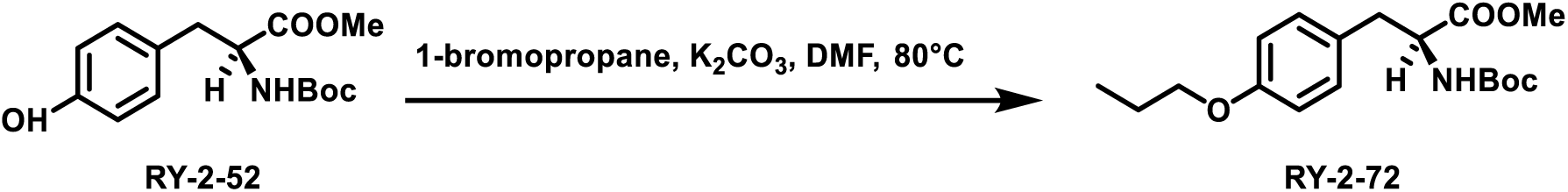

RY-2-52 (295 mg, 1 mmol, 1.0 eq) was dissolved in DMF (5 mL) followed by the addition of 1-bromopropane (246 mg, 2 mmol, 2.0 eq) and potassium carbonate (276 mg, 2 mmol, 2.0 eq). The reaction mixture was heated at 80°C and stirred overnight. Upon completion, the reaction mixture was quenched with water (20 mL) and extracted with ethyl acetate (25 mL x 3). The organic phase was washed with brine (20 mL x 3) and dried over sodium sulfate. The solvent was removed under reduced pressure and the residue was subjected to flash column chromatography (30% ethyl acetate in hexane). The product was obtained as white solid with a yield of 154 mg, 46%. ^1^H NMR (400 MHz, Chloroform-*d*) δ 7.02 (d, *J* = 8.5 Hz, 2H), 6.82 (d, *J* = 8.6 Hz, 2H), 4.95 (d, *J* = 8.3 Hz, 1H), 4.53 (d, *J* = 7.8 Hz, 1H), 3.89 (t, *J* = 6.6 Hz, 2H), 3.71 (s, 3H), 3.04 (dt, *J* = 13.9, 6.5 Hz, 2H), 1.79 (h, *J* = 7.1 Hz, 2H), 1.42 (s, 9H), 1.03 (t, *J* = 7.4 Hz, 3H).

#### Synthesis of RY-2-88

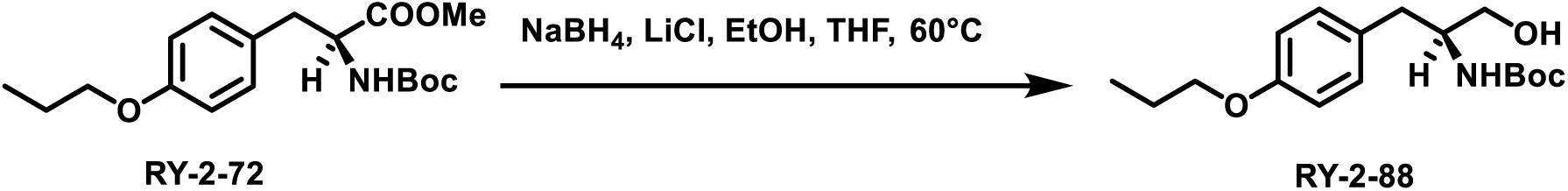

RY-2-72 (1000 mg, 3 mmol, 1.0 eq) was dissolved in ethanol (15 mL) and THF (15 mL) followed by the addition of sodium borohydride (454 mg, 12 mmol, 4.0 eq) and lithium chloride (509 mg, 12 mmol, 4.0 eq). The reaction mixture was heated at 60°C and stirred overnight. Upon completion, the reaction mixture was slowly quenched with water (50 mL) and extracted with ethyl acetate (100 mL x 3). The organic phase was washed with brine (100 mL x 3) and dried over sodium sulfate. The solvent was removed under reduced pressure and the residue was subjected to flash column chromatography (60% ethyl acetate in hexane). The product was obtained as white solid with a yield of 760 mg, 82%. ^1^H NMR (400 MHz, Chloroform-*d*) δ 7.11 (d, *J* = 8.6 Hz, 2H), 6.83 (d, *J* = 8.6 Hz, 2H), 4.72 (s, 1H), 3.90 (t, *J* = 6.6 Hz, 2H), 3.81 (s, 1H), 3.72 – 3.48 (m, 2H), 2.77 (d, *J* = 7.1 Hz, 2H), 1.80 (h, *J* = 7.1 Hz, 2H), 1.42 (s, 9H), 1.03 (t, *J* = 7.4 Hz, 3H).

#### Synthesis of XML-2

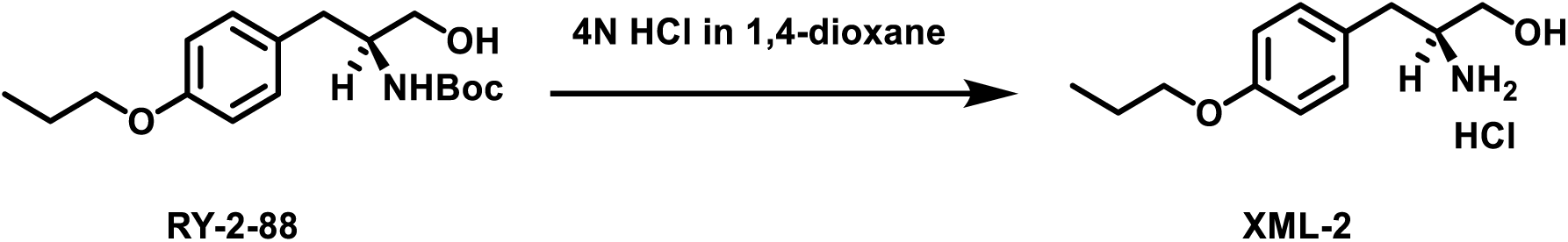

To a round-bottom flask was added RY-2-88 (300 mg, 1 mmol) and 4N HCl in 1,4-dioxane (5 mL). The reaction mixture was stirred for 2 h. Upon completion, the solvent was removed under reduced pressure. The residue was suspended in ethyl ether (5 mL) and filtered. The solid was washed with ethyl ether (10 mL) and dried under vacuum. The product was obtained as white solid with a yield of 210 mg, 88%. ^1^H NMR (400 MHz, Methanol-*d*4) δ 7.20 (d, *J* = 8.6 Hz, 2H), 6.92 (d, *J* = 8.6 Hz, 2H), 3.94 (t, *J* = 6.5 Hz, 2H), 3.71 (dd, *J* = 11.6, 3.6 Hz, 1H), 3.54 (dd, *J* = 11.6, 6.3 Hz, 1H), 3.47 – 3.36 (m, 1H), 2.89 (d, *J* = 7.6 Hz, 2H), 1.80 (h, *J* = 7.4 Hz, 2H), 1.05 (t, *J* = 7.4 Hz, 3H). ^13^C NMR (101 MHz, Methanol-*d*4) δ 158.50, 130.19, 127.36, 114.71, 69.20, 59.99, 54.61, 34.20, 22.26, 9.45. HRMS-ESI: calculated for [C12H20NO2]^+^: 210.14886, found: 210.14886. UPLC: tR = 3.617 min, >99% (A1/B1); tR = 2.798 min, >99% (A2/B2).

#### Synthesis of RY-2-74

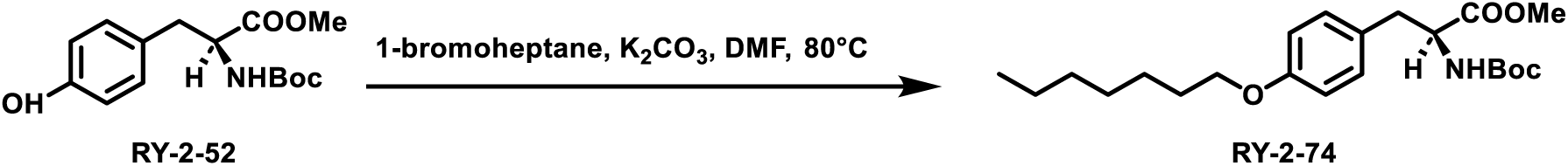

RY-2-52 (1181 mg, 4 mmol, 1.0 eq) was dissolved in DMF (20 mL) followed by the addition of 1-bromoheptane (1433 mg, 8 mmol, 2.0 eq) and potassium carbonate (1104 mg, 8 mmol, 2.0 eq). The reaction mixture was heated at 80°C and stirred overnight. Upon completion, the reaction mixture was quenched with water (100 mL) and extracted with ethyl acetate (100 mL x 3). The organic phase was washed with brine (100 mL x 3) and dried over sodium sulfate. The solvent was removed under reduced pressure and the residue was subjected to flash column chromatography (18% ethyl acetate in hexane). The product was obtained as white solid with a yield of 1.5 g, 95%. ^1^H NMR (400 MHz, Chloroform-*d*) δ 7.01 (d, *J* = 8.4 Hz, 2H), 6.81 (d, *J* = 8.6 Hz, 2H), 4.94 (d, *J* = 8.3 Hz, 1H), 4.53 (d, *J* = 6.9 Hz, 1H), 3.92 (t, *J* = 6.6 Hz, 2H), 3.71 (s, 3H), 3.15 – 2.84 (m, 2H), 1.87 – 1.68 (m, 2H), 1.57 (s, 1H), 1.52 – 1.17 (m, 17H), 1.00 – 0.79 (m, 3H).

#### Synthesis of RY-2-89

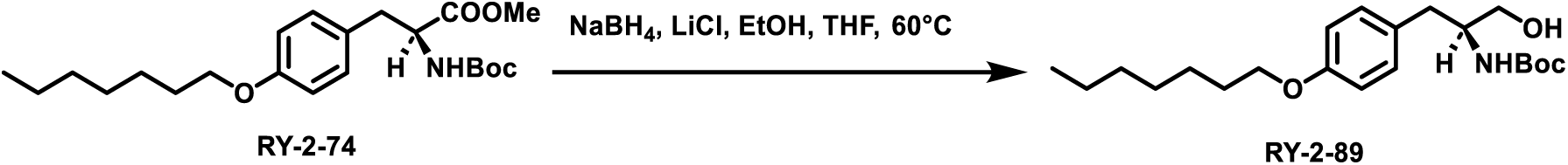

RY-2-74 (1000 mg, 2.5 mmol, 1.0 eq) was dissolved in ethanol (10 mL) and THF (10 mL) followed by the addition of sodium borohydride (378 mg, 10 mmol, 4.0 eq) and lithium chloride (424 mg, 10 mmol, 4.0 eq). The reaction mixture was heated at 60°C and stirred overnight. Upon completion, the reaction mixture was slowly quenched with water (50 mL) and extracted with ethyl acetate (100 mL x 3). The organic phase was washed with brine (100 mL x 3) and dried over sodium sulfate. The solvent was removed under reduced pressure and the residue was subjected to flash column chromatography (60% ethyl acetate in hexane). The product was obtained as white solid with a yield of 700 mg, 75%. ^1^H NMR (400 MHz, Chloroform-*d*) δ 7.10 (d, *J* = 8.5 Hz, 2H), 6.83 (d, *J* = 8.6 Hz, 2H), 4.74 (s, 1H), 3.93 (t, *J* = 6.6 Hz, 2H), 3.81 (s, 1H), 3.60 (ddd, *J* = 46.4, 11.0, 4.6 Hz, 2H), 2.77 (d, *J* = 7.1 Hz, 2H), 1.90 – 1.67 (m, 2H), 1.54 – 1.21 (m, 17H), 1.02 – 0.76 (m, 3H).

#### Synthesis of XML-3

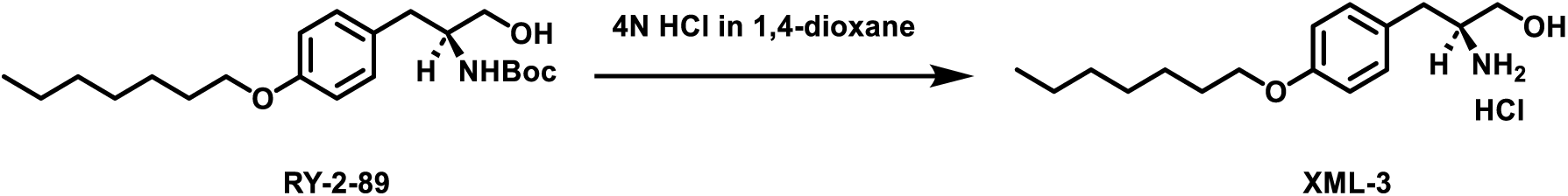

To a round-bottom flask was added RY-2-89 (300 mg, 0.8 mmol) and 4N HCl in 1,4-dioxane (5 mL). The reaction mixture was stirred for 2 h. Upon completion, the solvent was removed under reduced pressure. The residue was suspended in ethyl ether (5 mL) and filtered. The solid was washed with ethyl ether (10 mL) and dried under vacuum. The product was obtained as white solid with a yield of 207 mg, 84%. ^1^H NMR (400 MHz, Methanol-*d*4) δ 7.19 (d, *J* = 8.6 Hz, 2H), 6.91 (d, *J* = 8.7 Hz, 2H), 3.97 (t, *J* = 6.4 Hz, 2H), 3.71 (dd, *J* = 11.6, 3.5 Hz, 1H), 3.54 (dd, *J* = 11.6, 6.3 Hz, 1H), 3.47 – 3.36 (m, 1H), 2.98 – 2.80 (m, 2H), 1.87 – 1.69 (m, 2H), 1.57 – 1.25 (m, 8H), 1.03 – 0.85 (m, 3H). ^13^C NMR (101 MHz, Methanol-*d*4) δ 158.49, 129.96, 127.34, 114.57, 67.63, 59.99, 54.60, 34.20, 31.59, 29.02, 28.80, 25.75, 22.27, 13.03. HRMS-ESI: calculated for [C16H28NO2]^+^: 266.21146, found: 266.21115. UPLC: tR = 5.036 min, >99% (A1/B1); tR = 3.773 min, >99% (A2/B2).

#### Synthesis of RY-2-70

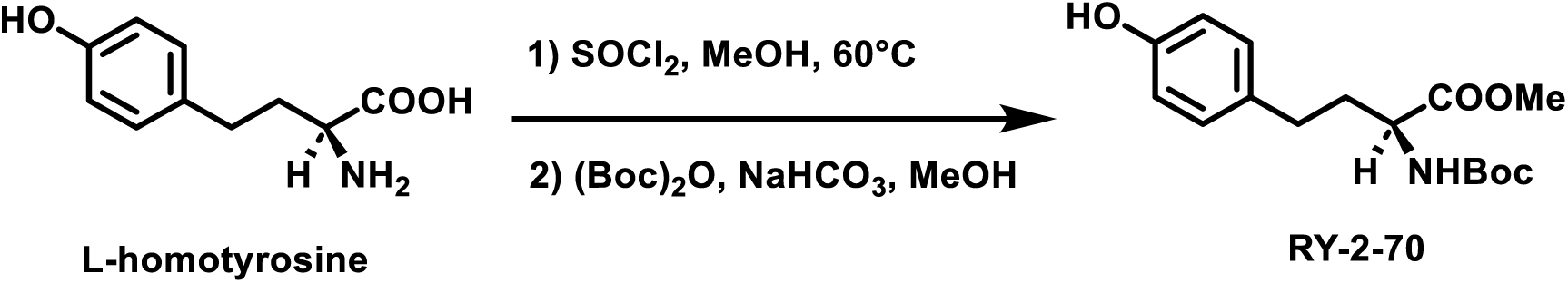

L-homotyrosine (2760 mg, 10 mmol, 1.0 eq) was dissolved in methanol (100 mL) followed by the addition of thionyl chloride (1190 mg, 100 mmol, 10.0 eq). The reaction mixture was stirred at 60°C overnight. Upon completion, the solvent was removed under reduced pressure and the residue was washed with ethyl ether (100 mL). The product was used without further purification. L-homotyrosine methyl ester (1045 mg, 5 mmol, 1.0 eq) was dissolved in methanol (40 mL) followed by the addition of sodium bicarbonate (840 mg, 10 mmol, 2.0 eq). A solution of Boc anhydride (1308 mg, 6 mmol, 1.2 eq) in methanol (10 mL) was added. The reaction was stirred at room temperature overnight. Upon completion, the solvent was dried under reduced pressure and the residue was re-dissolved in water (200 mL). The mixture was acidified with 1N HCl until pH = 4 and extracted with ethyl acetate (100 mL x 3). The organic phase was washed with brine (100 mL x 3) and dried over sodium sulfate. The solvent was removed under reduced pressure and the residue was subjected to flash column chromatography (35% ethyl acetate in hexane). The product was obtained as colorless oil with a yield of 860 mg, 56%. ^1^H NMR (400 MHz, Chloroform-*d*) δ 7.00 (d, *J* = 8.4 Hz, 2H), 6.75 (d, *J* = 8.4 Hz, 2H), 5.12 (d, *J* = 8.5 Hz, 1H), 4.48 – 4.22 (m, 1H), 3.71 (s, 3H), 2.58 (t, *J* = 8.0 Hz, 2H), 2.20 – 2.00 (m, 1H), 1.96 – 1.74 (m, 1H), 1.46 (s, 9H).

#### Synthesis of RY-2-76-1

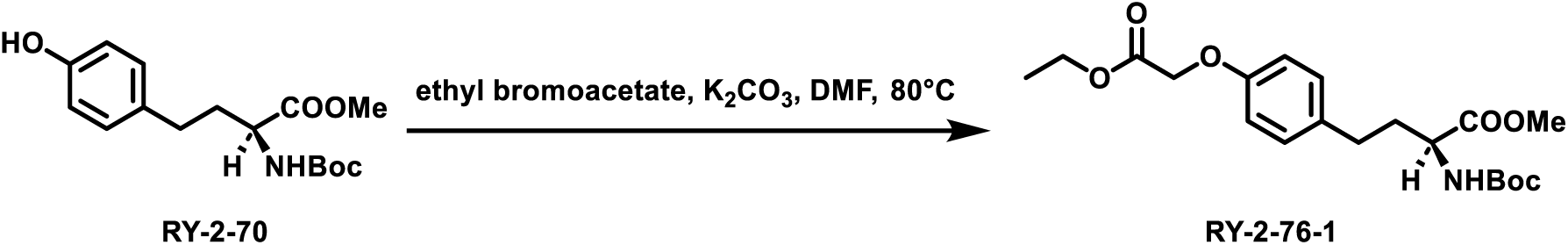

RY-2-70 (800 mg, 2.6 mmol, 1.0 eq) was dissolved in DMF (10 mL) followed by the addition of ethyl bromoacetate (868 mg, 5.2 mmol, 2.0 eq) and potassium carbonate (718 mg, 5.2 mmol, 2.0 eq). The reaction mixture was heated at 80°C and stirred overnight. Upon completion, the reaction mixture was quenched with water (50 mL) and extracted with ethyl acetate (50 mL x 3). The organic phase was washed with brine (50 mL x 3) and dried over sodium sulfate. The solvent was removed under reduced pressure and the residue was subjected to flash column chromatography (25% ethyl acetate in hexane). The product was obtained as light yellow oil with a yield of 1.0 g, 98%. ^1^H NMR (400 MHz, Chloroform-*d*) δ 7.09 (d, *J* = 8.5 Hz, 2H), 6.83 (d, *J* = 8.6 Hz, 2H), 5.08 (d, *J* = 8.5 Hz, 1H), 4.59 (s, 2H), 4.40 – 4.30 (m, 1H), 3.71 (s, 3H), 2.72 – 2.56 (m, 2H), 2.19 – 2.01 (m, 2H), 2.00 – 1.83 (m, 1H), 1.45 (s, 9H), 1.37 – 1.15 (m, 6H).

#### Synthesis of RY-2-79

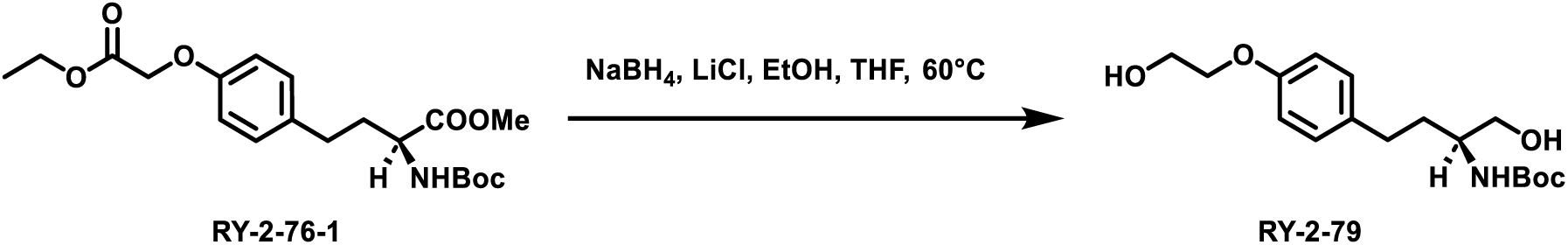

RY-2-76-1 (800 mg, 2 mmol, 1.0 eq) was dissolved in ethanol (10 mL) and THF (10 mL) followed by the addition of sodium borohydride (605 mg, 16 mmol, 8.0 eq) and lithium chloride (678 mg, 16 mmol, 8.0 eq). The reaction mixture was heated at 60°C and stirred overnight. Upon completion, the reaction mixture was slowly quenched with water (100 mL) and extracted with ethyl acetate (100 mL x 3). The organic phase was washed with brine (100 mL x 3) and dried over sodium sulfate. The solvent was removed under reduced pressure and the residue was subjected to flash column chromatography (90% ethyl acetate in hexane). The product was obtained as white solid with a yield of 246 mg, 38%. ^1^H NMR (400 MHz, Methanol-*d*4) δ 7.12 (d, *J* = 8.6 Hz, 2H), 6.87 (d, *J* = 8.6 Hz, 2H), 4.08 – 3.95 (m, 2H), 3.93 – 3.75 (m, 2H), 3.60 – 3.41 (m, 3H), 2.78 – 2.50 (m, 2H), 1.94 – 1.78 (m, 1H), 1.74 – 1.58 (m, 1H), 1.47 (s, 9H).

#### Synthesis of XML-4

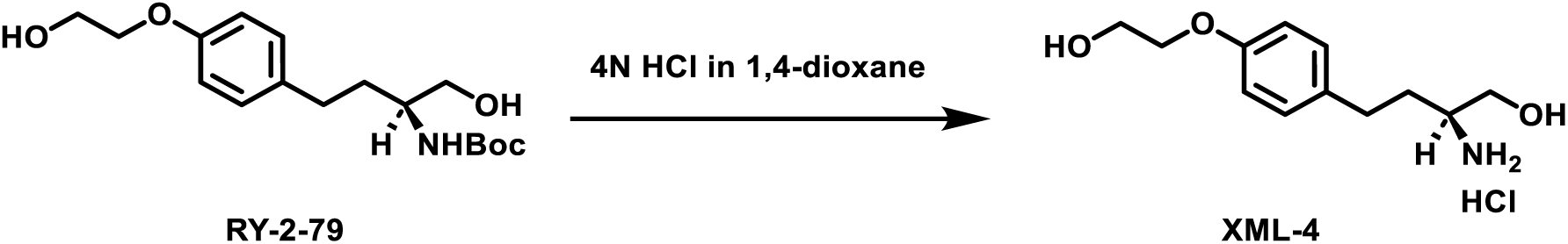

To a round-bottom flask was added RY-2-79 (200 mg, 0.6 mmol) and 4N HCl in 1,4-dioxane (4 mL). The reaction mixture was stirred for 2 h. Upon completion, the solvent was removed under reduced pressure. The residue was suspended in ethyl ether (5 mL) and filtered. The solid was washed with ethyl ether (10 mL) and dried under vacuum. The product was obtained as white solid with a yield of 154 mg, 96%. ^1^H NMR (400 MHz, Methanol-*d*4) δ 7.17 (d, *J* = 8.6 Hz, 2H), 6.91 (d, *J* = 8.6 Hz, 2H), 4.10 – 3.95 (m, 2H), 3.93 – 3.83 (m, 2H), 3.80 (dd, *J* = 11.8, 3.6 Hz, 1H), 3.62 (dd, J = 11.7, 6.7 Hz, 1H), 3.28 – 3.13 (m, 1H), 2.79 – 2.60 (m, 2H), 2.05 – 1.79 (m, 2H). ^13^C NMR (101 MHz, Methanol-*d4*) δ 157.62, 132.57, 128.94, 114.40, 69.19, 60.53, 60.34, 52.76, 31.12, 30.21. HRMS-ESI: calculated for [C12H20NO3]^+^: 226.14377, found: 226.14360. UPLC: tR = 2.459 min, 80.5% (A1/B1); tR = 2.074 min, >99% (A2/B2).

#### Synthesis of RY-2-76-2

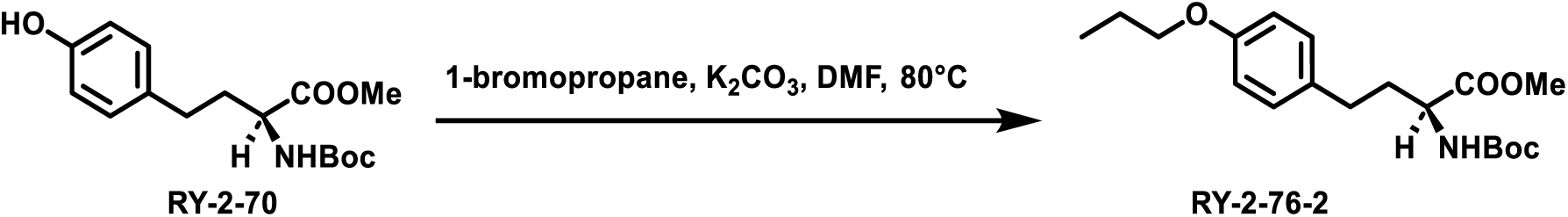

RY-2-70 (800 mg, 2.6 mmol, 1.0 eq) was dissolved in DMF (10 mL) followed by the addition of 1-bromopropane (640 mg, 5.2 mmol, 2.0 eq) and potassium carbonate (718 mg, 5.2 mmol, 2.0 eq). The reaction mixture was heated at 80°C and stirred overnight. Upon completion, the reaction mixture was quenched with water (50 mL) and extracted with ethyl acetate (50 mL x 3). The organic phase was washed with brine (50 mL x 3) and dried over sodium sulfate. The solvent was removed under reduced pressure and the residue was subjected to flash column chromatography (30% ethyl acetate in hexane). The product was obtained as white solid with a yield of 870 mg, 95%. ^1^H NMR (400 MHz, Chloroform-*d*) δ 7.07 (d, *J* = 8.5 Hz, 2H), 6.82 (d, *J* = 8.6 Hz, 2H), 5.05 (d, *J* = 8.6 Hz, 1H), 4.34 (q, *J* = 7.3 Hz, 1H), 3.89 (t, *J* = 6.6 Hz, 2H), 3.71 (s, 3H), 2.75 – 2.46 (m, 2H), 2.18 – 2.06 (m, 1H), 1.97 – 1.85 (m, 1H), 1.84 – 1.73 (m, 2H), 1.45 (s, 9H), 1.02 (t, *J* = 7.4 Hz, 3H).

#### Synthesis of RY-2-80

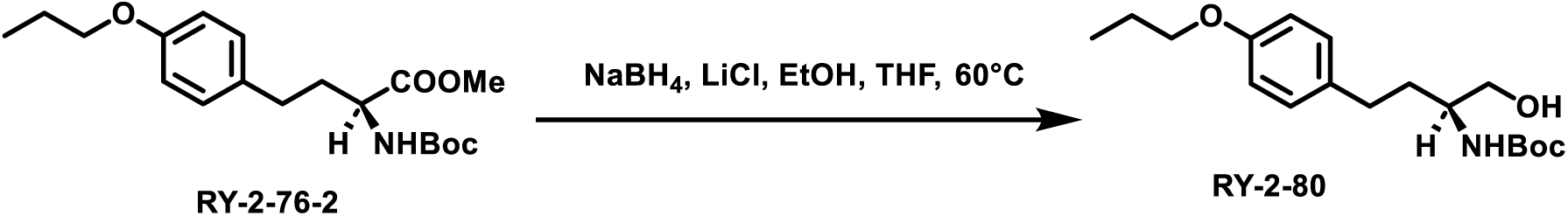

RY-2-76-2 (700 mg, 2 mmol, 1.0 eq) was dissolved in ethanol (10 mL) and THF (10 mL) followed by the addition of sodium borohydride (303 mg, 8 mmol, 4.0 eq) and lithium chloride (339 mg, 8 mmol, 4.0 eq). The reaction mixture was heated at 60°C and stirred overnight. Upon completion, the reaction mixture was slowly quenched with water (100 mL) and extracted with ethyl acetate (100 mL x 3). The organic phase was washed with brine (100 mL x 3) and dried over sodium sulfate. The solvent was removed under reduced pressure and the residue was subjected to flash column chromatography (50% ethyl acetate in hexane). The product was obtained as white solid with a yield of 483 mg, 75%. ^1^H NMR (400 MHz, Chloroform-*d*) δ 7.08 (d, *J* = 8.6 Hz, 2H), 6.81 (d, *J* = 8.6 Hz, 2H), 4.68 (s, 1H), 3.89 (t, *J* = 6.6 Hz, 2H), 3.76 – 3.46 (m, 3H), 2.77 – 2.48 (m, 2H), 2.30 (s, 1H), 1.92 – 1.63 (m, 4H), 1.45 (s, 9H), 1.02 (t, *J* = 7.4 Hz, 3H).

#### Synthesis of XML-5

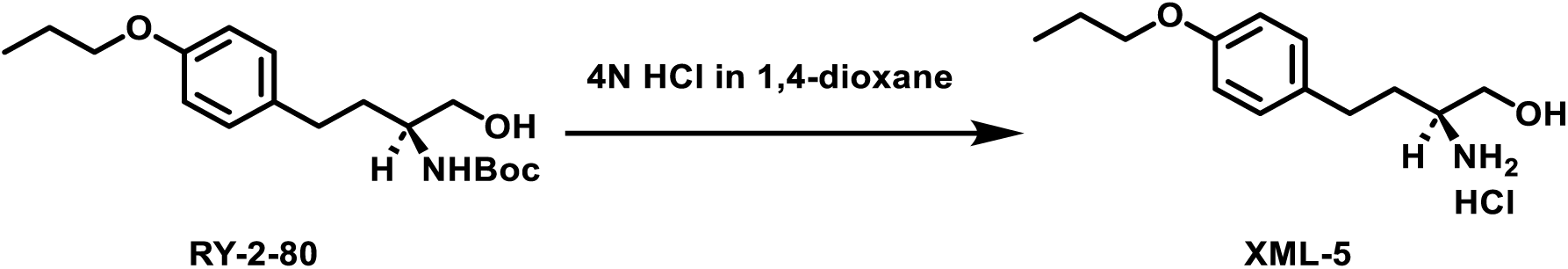

To a round-bottom flask was added RY-2-80 (200 mg, 0.6 mmol) and 4N HCl in 1,4-dioxane (4 mL). The reaction mixture was stirred for 2 h. Upon completion, the solvent was removed under reduced pressure. The residue was suspended in ethyl ether (5 mL) and filtered. The solid was washed with ethyl ether (10 mL) and dried under vacuum. The product was obtained as white solid with a yield of 154 mg, 97%. ^1^H NMR (400 MHz, Methanol-*d*4) δ 7.15 (d, *J* = 8.6 Hz, 2H), 6.86 (d, *J* = 8.6 Hz, 2H), 3.92 (t, *J* = 6.5 Hz, 2H), 3.81 (dd, *J* = 11.7, 3.6 Hz, 1H), 3.62 (dd, *J* = 11.7, 6.7 Hz, 1H), 3.27 – 3.10 (m, 1H), 2.79 – 2.58 (m, 2H), 2.03 – 1.84 (m, 2H), 1.84 – 1.70 (m, 2H), 1.05 (t, *J* = 7.4 Hz, 3H). ^13^C NMR (101 MHz, Methanol-*d*4) δ 157.79, 132.23, 128.89, 114.31, 69.20, 60.54, 52.77, 31.14, 30.22, 22.30, 9.46. HRMS-ESI: calculated for [C13H22NO2]^+^: 224.16451, found: 224.16443. UPLC: tR = 3.870 min, >99% (A1/B1); tR = 2.977 min, >99% (A2/B2).

#### Synthesis of RY-2-76-3

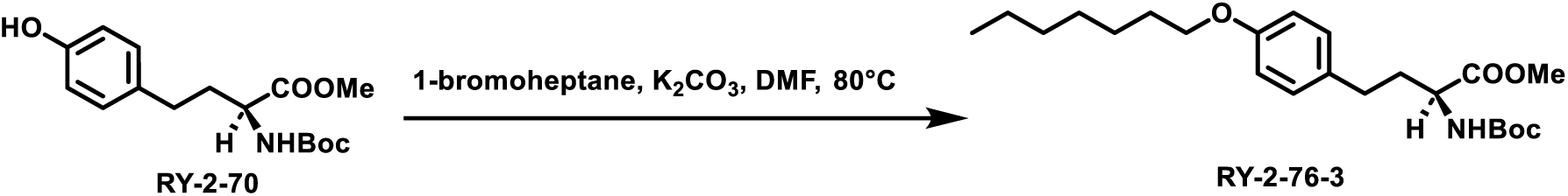

RY-2-70 (800 mg, 2.6 mmol, 1.0 eq) was dissolved in DMF (10 mL) followed by the addition of 1-bromoheptane (931 mg, 5.2 mmol, 2.0 eq) and potassium carbonate (718 mg, 5.2 mmol, 2.0 eq). The reaction mixture was heated at 80°C and stirred overnight. Upon completion, the reaction mixture was quenched with water (50 mL) and extracted with ethyl acetate (50 mL x 3). The organic phase was washed with brine (50 mL x 3) and dried over sodium sulfate. The solvent was removed under reduced pressure and the residue was subjected to flash column chromatography (30% ethyl acetate in hexane). The product was obtained as white solid with a yield of 930 mg, 88%. ^1^H NMR (400 MHz, Chloroform-*d*) δ 7.07 (d, *J* = 8.6 Hz, 2H), 6.81 (d, *J* = 8.5 Hz, 2H), 5.04 (d, *J* = 8.6 Hz, 1H), 4.34 (d, *J* = 7.1 Hz, 1H), 3.92 (t, *J* = 6.6 Hz, 2H), 3.71 (s, 3H), 2.72 – 2.49 (m, 2H), 2.22 – 2.00 (m, 1H), 1.99 – 1.83 (m, 1H), 1.83 – 1.70 (m, 2H), 1.53 – 1.18 (m, 17H), 1.03 – 0.78 (m, 3H).

#### Synthesis of RY-2-81

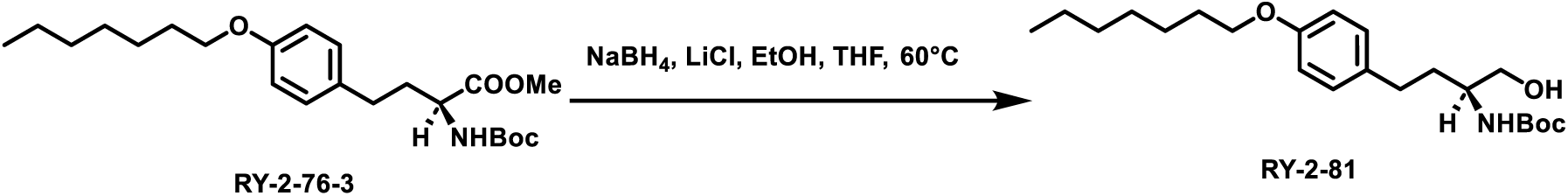

RY-2-76-3 (800 mg, 2 mmol, 1.0 eq) was dissolved in ethanol (10 mL) and THF (10 mL) followed by the addition of sodium borohydride (303 mg, 8 mmol, 4.0 eq) and lithium chloride (339 mg, 8 mmol, 4.0 eq). The reaction mixture was heated at 60°C and stirred overnight. Upon completion, the reaction mixture was slowly quenched with water (100 mL) and extracted with ethyl acetate (100 mL x 3). The organic phase was washed with brine (100 mL x 3) and dried over sodium sulfate. The solvent was removed under reduced pressure and the residue was subjected to flash column chromatography (40% ethyl acetate in hexane). The product was obtained as white solid with a yield of 485 mg, 64%. ^1^H NMR (400 MHz, Chloroform-*d*) δ 7.08 (d, *J* = 8.3 Hz, 2H), 6.81 (d, *J* = 7.5 Hz, 2H), 4.65 (s, 1H), 4.00 – 3.81 (m, 2H), 3.77 – 3.41 (m, 3H), 2.79 – 2.49 (m, 2H), 2.08 (s, 1H), 1.87 – 1.63 (m, 4H), 1.55 – 1.20 (m, 17H), 0.98 – 0.72 (m, 3H).

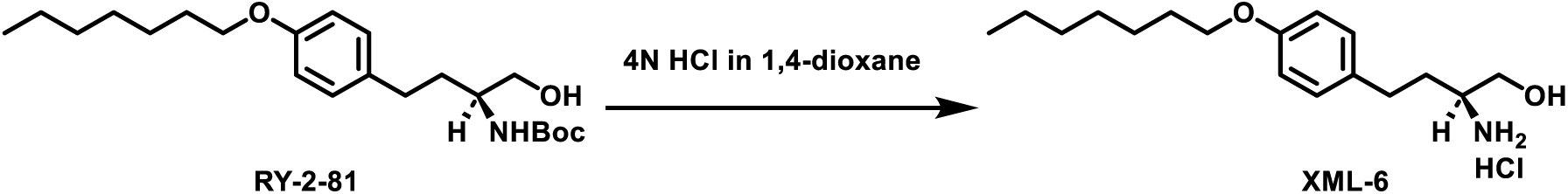

#### Synthesis of XML-6

To a round-bottom flask was added RY-2-81 (300 mg, 0.6 mmol) and 4N HCl in 1,4-dioxane (4 mL). The reaction mixture was stirred for 2 h. Upon completion, the solvent was removed under reduced pressure. The residue was suspended in ethyl ether (5 mL) and filtered. The solid was washed with ethyl ether (10 mL) and dried under vacuum. The product was obtained as white solid with a yield of 211 mg, 85%. ^1^H NMR (400 MHz, Methanol-*d*4) δ 7.15 (d, *J* = 8.5 Hz, 2H), 6.86 (d, *J* = 8.5 Hz, 2H), 3.95 (t, *J* = 6.4 Hz, 2H), 3.80 (dd, *J* = 11.7, 3.2 Hz, 1H), 3.61 (dd, *J* = 11.7, 6.4 Hz, 1H), 3.20 (s, 1H), 2.81 – 2.58 (m, 2H), 2.04 – 1.82 (m, 2H), 1.82 – 1.67 (m, 2H), 1.57 – 1.24 (m, 8H), 0.93 (t, *J* = 6.8 Hz, 3H). ^13^C NMR (101 MHz, Methanol-*d*4) δ 157.79, 132.19, 128.88, 114.29, 67.62, 60.53, 52.74, 31.59, 31.14, 30.21, 29.06, 28.82, 25.76, 22.28, 13.03. HRMS-ESI: calculated for [C17H30NO2]^+^: 280.22711, found: 280.22657. UPLC: tR = 5.142 min, >99% (A1/B1); tR = 3.896 min, >99% (A2/B2).

#### Synthesis of RY-2-68

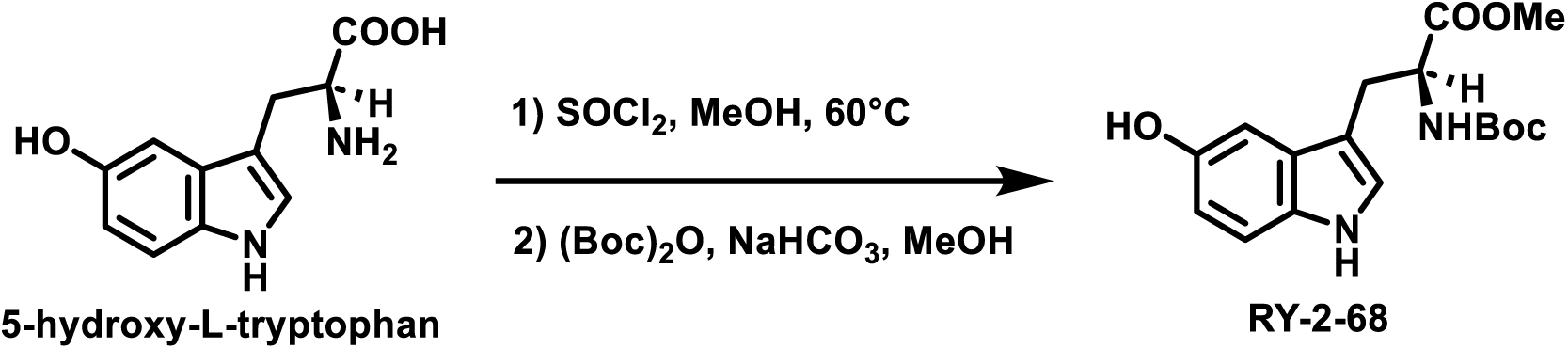

5-Hydroxy-L-tryptophan (2200 mg, 10 mmol, 1.0 eq) was dissolved in methanol (100 mL) followed by the addition of thionyl chloride (1190 mg, 100 mmol, 10.0 eq). The reaction mixture was stirred at 60°C overnight. Upon completion, the solvent was removed under reduced pressure and the residue was washed with ethyl ether (100 mL). The product was used without further purification. 5-Hydroxy-L-tryptophan methyl ester (1638 mg, 7 mmol, 1.0 eq) was dissolved in methanol (50 mL) followed by the addition of sodium bicarbonate (1176 mg, 14 mmol, 2.0 eq). A solution of Boc anhydride (1833 mg, 8.4 mmol, 1.2 eq) in methanol (10 mL) was added. The reaction was stirred at room temperature overnight. Upon completion, the solvent was dried under reduced pressure and the residue was re-dissolved in water (200 mL). The mixture was acidified with 1N HCl until pH = 4 and extracted with ethyl acetate (100 mL x 3). The organic phase was washed with brine (100 mL x 3) and dried over sodium sulfate. The solvent was removed under reduced pressure and the residue was subjected to flash column chromatography (80% ethyl acetate in hexane). The product was obtained as brown solid with a yield of 1050 mg, 45%. ^1^H NMR (400 MHz, Methanol-*d*4) δ 7.17 (d, *J* = 8.7 Hz, 1H), 7.02 (s, 1H), 6.92 (d, *J* = 2.3 Hz, 1H), 6.68 (dd, *J* = 8.7, 2.3 Hz, 1H), 4.50 – 4.29 (m, 1H), 3.68 (s, 3H), 3.27 – 2.91 (m, 2H), 1.41 (s, 9H).

#### Synthesis of RY-2-71

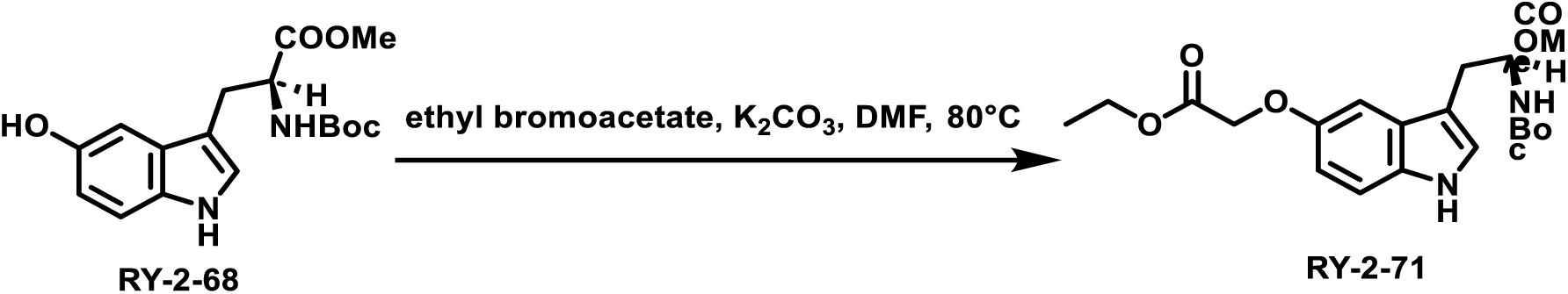

RY-2-68 (1002 mg, 3 mmol, 1.0 eq) was dissolved in DMF (20 mL) followed by the addition of ethyl bromoacetate (1002 mg, 6 mmol, 2.0 eq) and potassium carbonate (828 mg, 6 mmol, 2.0 eq). The reaction mixture was heated at 80°C and stirred overnight. Upon completion, the reaction mixture was quenched with water (100 mL) and extracted with ethyl acetate (100 mL x 3). The organic phase was washed with brine (100 mL x 3) and dried over sodium sulfate. The solvent was removed under reduced pressure and the residue was subjected to flash column chromatography (50% ethyl acetate in hexane). The product was obtained as light-yellow oil with a yield of 930 mg, 74%. ^1^H NMR (300 MHz, Chloroform-*d*) δ 8.26 (s, 1H), 7.38 – 7.15 (m, 1H), 7.08 – 6.80 (m, 3H), 5.10 (d, *J* = 8.4 Hz, 1H), 4.66 (s, 3H), 4.29 (q, *J* = 7.1 Hz, 2H), 3.67 (s, 3H), 3.22 (d, *J* = 5.6 Hz, 2H), 1.43 (s, 9H), 1.26 (t, *J* = 7.2 Hz, 3H).

#### Synthesis of RY-2-75

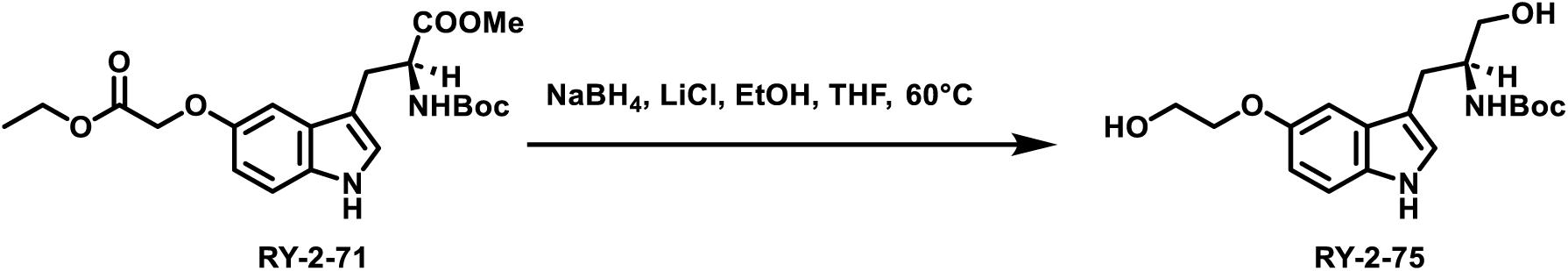

RY-2-71 (800 mg, 2 mmol, 1.0 eq) was dissolved in ethanol (10 mL) and THF (10 mL) followed by the addition of sodium borohydride (605 mg, 16 mmol, 8.0 eq) and lithium chloride (678 mg, 16 mmol, 8.0 eq). The reaction mixture was heated at 60°C and stirred overnight. Upon completion, the reaction mixture was slowly quenched with water (100 mL) and extracted with ethyl acetate (100 mL x 3). The organic phase was washed with brine (100 mL x 3) and dried over sodium sulfate. The solvent was removed under reduced pressure and the residue was subjected to flash column chromatography (90% ethyl acetate in hexane). The product was obtained as brown solid with a yield of 320 mg, 48%. ^1^H NMR (300 MHz, Chloroform-*d*) δ 8.07 (s, 1H), 7.23 (s, 1H), 7.17 (d, *J* = 2.4 Hz, 1H), 7.02 (s, 1H), 6.88 (dd, *J* = 8.8, 2.4 Hz, 1H), 4.86 (s, 1H), 4.29 – 4.07 (m, 2H), 4.05 – 3.81 (m, 2H), 3.63 (qd, *J* = 10.9, 4.7 Hz, 2H), 2.94 (d, *J* = 6.8 Hz, 2H), 2.34 (s, 2H), 1.42 (s, 9H).

#### Synthesis of XML-7

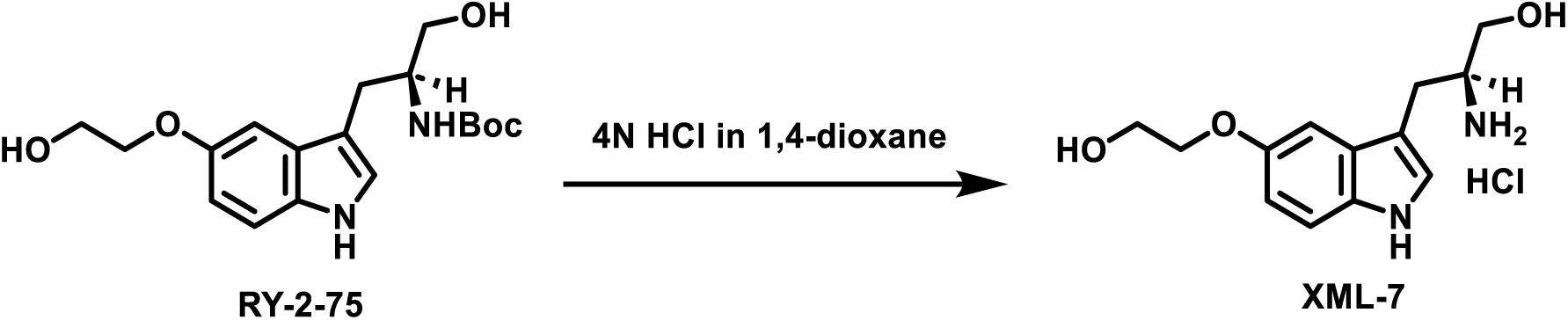

To a round-bottom flask was added RY-2-75 (300 mg, 0.9 mmol) and 4N HCl in 1,4-dioxane (5 mL). The reaction mixture was stirred for 2 h. Upon completion, the solvent was removed under reduced pressure. The residue was suspended in ethyl ether (5 mL) and filtered. The solid was washed with ethyl ether (10 mL) and dried under vacuum. The product was obtained as yellow solid with a yield of 253 mg, 88%. ^1^H NMR (400 MHz, Methanol-*d*4) δ 7.29 (d, *J* = 8.8 Hz, 1H), 7.18 (s, 1H), 7.15 (d, *J* = 2.3 Hz, 1H), 6.87 (dd, *J* = 8.8, 2.3 Hz, 1H), 4.11 (t, *J* = 4.8 Hz, 2H), 3.91 (t, *J* = 5.0 Hz, 3H), 3.79 (dd, *J* = 11.5, 3.6 Hz, 1H), 3.63 (dd, *J* = 11.5, 6.5 Hz, 1H), 3.57 – 3.50 (m, 1H), 3.16 – 3.00 (m, 2H). ^13^C NMR (101 MHz, Methanol-*d*4) δ 153.13, 132.27, 127.40, 124.35, 112.24, 111.86, 107.83, 101.10, 70.11, 60.59, 53.67, 24.97. MS (ESI) calculated for [C13H19N2O3]^+^: 251.3, found: 251.2. UPLC: tR = 2.440 min, >99% (A1/B1); tR = 2.080 min, >99% (A2/B2).

#### Synthesis of RY-2-90-1

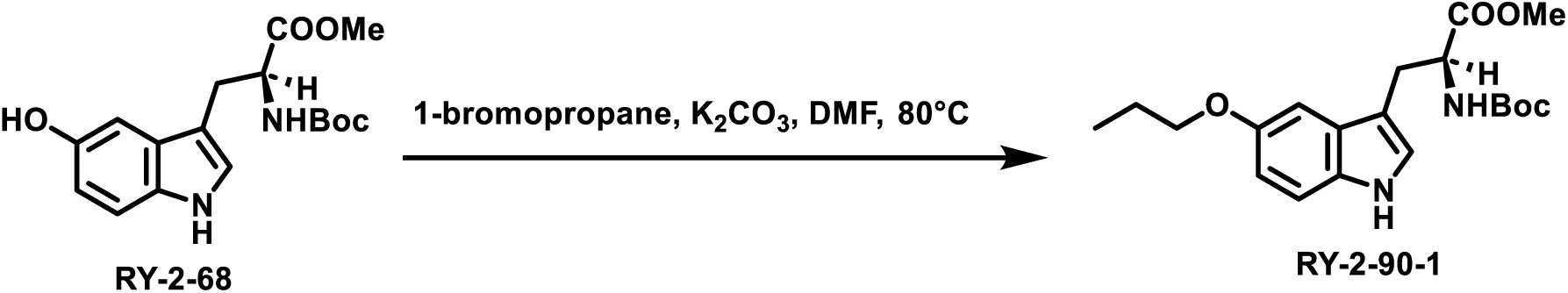

RY-2-68 (1002 mg, 3 mmol, 1.0 eq) was dissolved in DMF (15 mL) followed by the addition of 1-bromopropane (756 mg, 6 mmol, 2.0 eq) and potassium carbonate (828 mg, 6 mmol, 2.0 eq). The reaction mixture was heated at 80°C and stirred overnight. Upon completion, the reaction mixture was quenched with water (100 mL) and extracted with ethyl acetate (100 mL x 3). The organic phase was washed with brine (100 mL x 3) and dried over sodium sulfate. The solvent was removed under reduced pressure and the residue was subjected to flash column chromatography (30% ethyl acetate in hexane). The product was obtained as orange solid with a yield of 380 mg, 34%. ^1^H NMR (400 MHz, Chloroform-*d*) δ 7.98 (s, 1H), 7.22 (d, *J* = 8.7 Hz, 1H), 6.99 (dd, *J* = 16.6, 2.3 Hz, 2H), 6.86 (dd, *J* = 8.8, 2.4 Hz, 1H), 5.09 (d, *J* = 8.4 Hz, 1H), 4.64 (d, *J* = 7.1 Hz, 1H), 3.97 (t, *J* = 6.6 Hz, 2H), 3.68 (s, 3H), 3.24 (d, *J* = 5.5 Hz, 2H), 1.83 (h, *J* = 7.1 Hz, 2H), 1.42 (s, 9H), 1.06 (t, *J* = 7.4 Hz, 3H).

#### Synthesis of RY-2-104

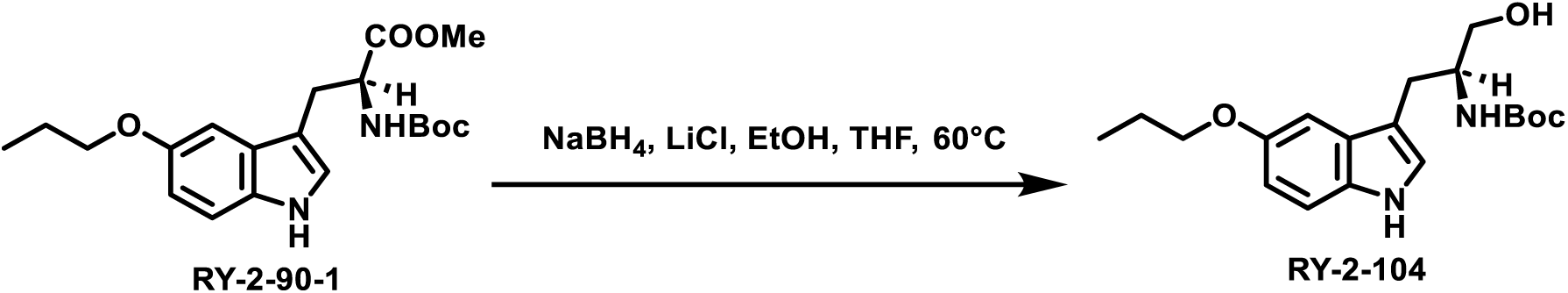

RY-2-9-1 (752 mg, 2 mmol, 1.0 eq) was dissolved in ethanol (10 mL) and THF (10 mL) followed by the addition of sodium borohydride (303 mg, 8 mmol, 4.0 eq) and lithium chloride (339 mg, 8 mmol, 4.0 eq). The reaction mixture was heated at 60°C and stirred overnight. Upon completion, the reaction mixture was slowly quenched with water (100 mL) and extracted with ethyl acetate (100 mL x 3). The organic phase was washed with brine (100 mL x 3) and dried over sodium sulfate. The solvent was removed under reduced pressure and the residue was subjected to flash column chromatography (70% ethyl acetate in hexane). The product was obtained as white solid with a yield of 670 mg, 96%. ^1^H NMR (300 MHz, Chloroform-*d*) δ 8.12 (s, 1H), 7.23 (d, *J* = 8.8 Hz, 1H), 7.16 – 6.95 (m, 2H), 6.86 (dd, *J* = 8.8, 2.4 Hz, 1H), 4.87 (d, *J* = 7.6 Hz, 1H), 3.97 (t, *J* = 6.6 Hz, 3H), 3.78 – 3.50 (m, 2H), 2.94 (d, *J* = 6.7 Hz, 2H), 2.51 (s, 1H), 1.83 (h, *J* = 7.1 Hz, 2H), 1.41 (s, 9H), 1.06 (t, *J* = 7.4 Hz, 3H).

#### Synthesis of XML-8

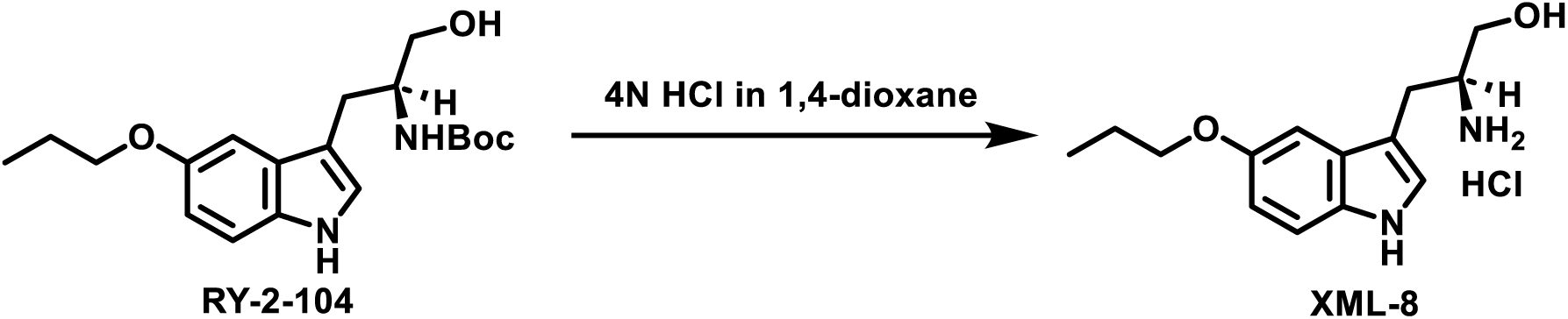

To a round-bottom flask was added RY-2-104 (600 mg, 1.7 mmol) and 4N HCl in 1,4-dioxane (5 mL). The reaction mixture was stirred for 2 h. Upon completion, the solvent was removed under reduced pressure. The residue was suspended in ethyl ether (5 mL) and filtered. The solid was washed with ethyl ether (10 mL) and dried under vacuum. The product was obtained as brown solid with a yield of 320 mg, 65%. ^1^H NMR (400 MHz, Methanol-*d*4) δ 7.29 (d, *J* = 8.8 Hz, 1H), 7.18 (s, 1H), 7.11 (d, *J* = 2.3 Hz, 1H), 6.82 (dd, *J* = 8.8, 2.4 Hz, 1H), 3.99 (t, *J* = 6.5 Hz, 2H), 3.79 (dd, *J* = 11.5, 3.7 Hz, 1H), 3.63 (dd, *J* = 11.5, 6.4 Hz, 1H), 3.58 – 3.48 (m, 1H), 3.18 – 2.98 (m, 2H), 1.93 – 1.74 (m, 2H), 1.09 (t, *J* = 7.4 Hz, 3H). ^13^C NMR (101 MHz, Methanol-*d*4) δ 153.28, 132.15, 127.41, 124.27, 112.16, 111.81, 107.77, 100.94, 70.18, 60.59, 53.67, 24.98, 22.53, 9.60. HRMS-ESI: calculated for [C14H21N2O2]^+^: 249.15975, found: 249.15944. UPLC: tR = 3.640 min, 90.03% (A1/B1); tR = 2.847 min, 86.85% (A2/B2).

#### Synthesis of RY-2-90-2

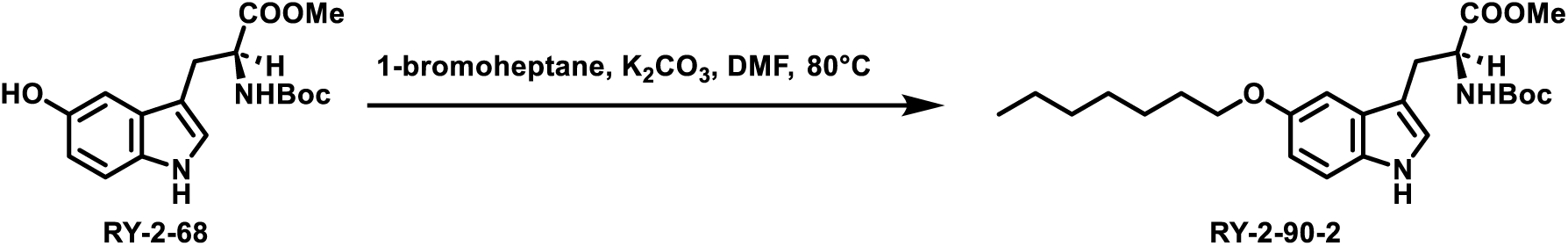

RY-2-68 (1002 mg, 3 mmol, 1.0 eq) was dissolved in DMF (15 mL) followed by the addition of 1-bromoheptane (1075 mg, 6 mmol, 2.0 eq) and cesium carbonate (828 mg, 6 mmol, 2.0 eq). The reaction mixture was heated at 80°C and stirred overnight. Upon completion, the reaction mixture was quenched with water (100 mL) and extracted with ethyl acetate (100 mL x 3). The organic phase was washed with brine (100 mL x 3) and dried over sodium sulfate. The solvent was removed under reduced pressure and the residue was subjected to flash column chromatography (30% ethyl acetate in hexane). The product was obtained as orange oil with a yield of 820 mg, 63%. ^1^H NMR (400 MHz, Chloroform-*d*) δ 8.06 (s, 1H), 7.22 (d, *J* = 8.7 Hz, 1H), 6.98 (dd, *J* = 16.3, 2.1 Hz, 2H), 6.85 (dd, *J* = 8.8, 2.4 Hz, 1H), 5.10 (d, *J* = 8.3 Hz, 1H), 4.74 – 4.55 (m, 1H), 4.00 (t, *J* = 6.6 Hz, 2H), 3.68 (s, 3H), 3.23 (d, *J* = 5.5 Hz, 2H), 1.88 – 1.74 (m, 2H), 1.67 – 1.27 (m, 22H), 1.00 – 0.80 (m, 3H).

#### Synthesis of RY-2-91-2

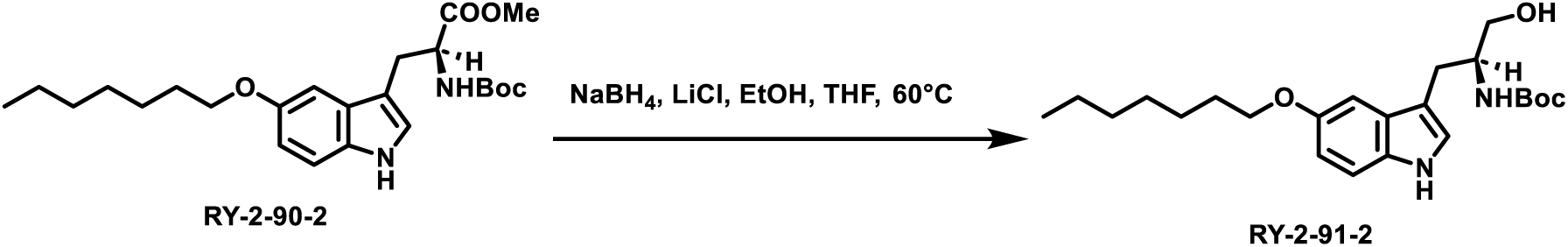

RY-2-90-2 (648 mg, 1.5 mmol, 1.0 eq) was dissolved in ethanol (15 mL) and THF (15 mL) followed by the addition of sodium borohydride (227 mg, 6 mmol, 4.0 eq) and lithium chloride (255 mg, 6 mmol, 4.0 eq). The reaction mixture was heated at 60°C and stirred overnight. Upon completion, the reaction mixture was slowly quenched with water (50 mL) and extracted with ethyl acetate (100 mL x 3). The organic phase was washed with brine (100 mL x 3) and dried over sodium sulfate. The solvent was removed under reduced pressure and the residue was subjected to flash column chromatography (60% ethyl acetate in hexane). The product was obtained as brown solid with a yield of 378 mg, 62%. ^1^H NMR (400 MHz, Chloroform-*d*) δ 8.04 (s, 1H), 7.23 (d, *J* = 8.8 Hz, 1H), 7.15 – 6.95 (m, 2H), 6.86 (dd, *J* = 8.7, 2.4 Hz, 1H), 4.85 (s, 1H), 4.00 (t, *J* = 6.6 Hz, 3H), 3.78 – 3.52 (m, 2H), 3.06 – 2.60 (m, 3H), 1.80 (p, *J* = 6.8 Hz, 2H), 1.63 – 1.22 (m, 20H), 0.99 – 0.80 (m, 3H).

#### Synthesis of XML-9

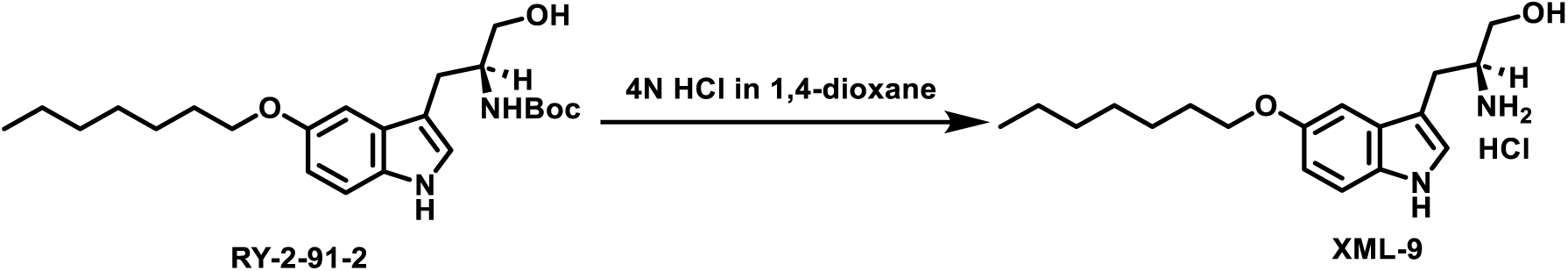

To a round-bottom flask was added RY-2-91-2 (300 mg, 0.7 mmol) and 4N HCl in 1,4-dioxane (5 mL). The reaction mixture was stirred for 2 h. Upon completion, the solvent was removed under reduced pressure. The residue was suspended in ethyl ether (5 mL) and filtered. The solid was washed with ethyl ether (10 mL) and dried under vacuum. The product was obtained as brown solid with a yield of 147 mg, 58%. ^1^H NMR (400 MHz, Methanol-*d*4) δ 7.28 (d, *J* = 8.8 Hz, 1H), 7.16 (s, 1H), 7.09 (d, *J* = 2.3 Hz, 1H), 6.83 (dd, *J* = 8.8, 2.4 Hz, 1H), 4.03 (t, *J* = 6.5 Hz, 2H), 3.79 (dd, *J* = 11.4, 3.7 Hz, 1H), 3.63 (dd, *J* = 11.5, 6.5 Hz, 1H), 3.58 – 3.46 (m, 1H), 3.19 – 2.96 (m, 2H), 1.91 – 1.68 (m, 2H), 1.63 – 1.20 (m, 9H), 1.07 – 0.79 (m, 3H). ^13^C NMR (101 MHz, Methanol-*d*4) δ 153.30, 127.42, 124.29, 112.16, 111.82, 107.77, 100.97, 68.63, 60.61, 53.69, 31.64, 29.35, 28.90, 25.90, 25.00, 22.30, 13.04. HRMS-ESI: calculated for [C18H29N2O2]^+^: 305.22235, found: 305.22212. UPLC: 4.962 min, 89.96% (A1/B1); tR = 3.743 min, 92.39% (A2/B2).

#### Synthesis of RY-2-110

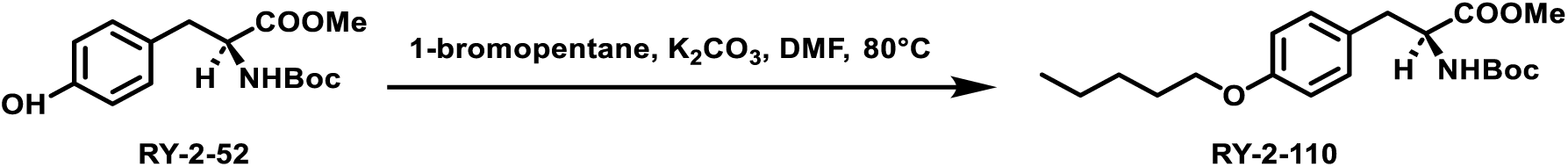

RY-2-52 (885 mg, 3 mmol, 1.0 eq) was dissolved in DMF (10 mL) followed by the addition of 1-bromopentane (906 mg, 6 mmol, 2.0 eq) and potassium carbonate (828 mg, 6 mmol, 2.0 eq). The reaction mixture was heated at 80°C and stirred overnight. Upon completion, the reaction mixture was quenched with water (50 mL) and extracted with ethyl acetate (50 mL x 3). The organic phase was washed with brine (50 mL x 3) and dried over sodium sulfate. The solvent was removed under reduced pressure and the residue was subjected to flash column chromatography (25% ethyl acetate in hexane). The product was obtained as white solid with a yield of 900 mg, 85%. ^1^H NMR (400 MHz, Chloroform-*d*) δ 7.01 (d, *J* = 8.5 Hz, 2H), 6.81 (d, *J* = 8.6 Hz, 2H), 4.95 (d, *J* = 8.3 Hz, 1H), 4.53 (d, *J* = 7.2 Hz, 1H), 3.92 (t, *J* = 6.6 Hz, 2H), 3.71 (s, 3H), 3.13 – 2.93 (m, 2H), 1.86 – 1.68 (m, 2H), 1.49 – 1.27 (m, 13H), 0.93 (t, *J* = 7.0 Hz, 3H).

#### Synthesis of RY-2-114

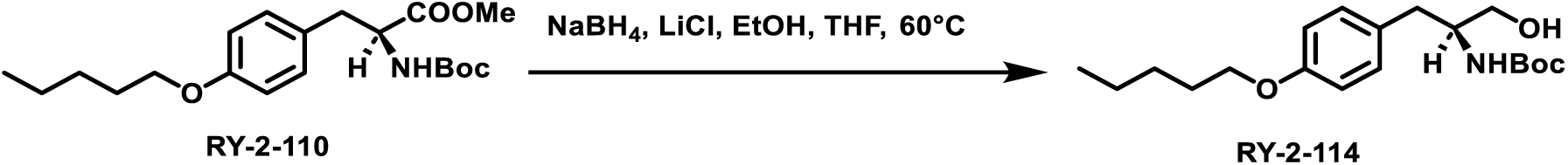

RY-2-110 (878 mg, 2.5 mmol, 1.0 eq) was dissolved in ethanol (10 mL) and THF (10 mL) followed by the addition of sodium borohydride (378 mg, 10 mmol, 4.0 eq) and lithium chloride (424 mg, 10 mmol, 4.0 eq). The reaction mixture was heated at 60°C and stirred overnight. Upon completion, the reaction mixture was slowly quenched with water (50 mL) and extracted with ethyl acetate (100 mL x 3). The organic phase was washed with brine (100 mL x 3) and dried over sodium sulfate. The solvent was removed under reduced pressure and the residue was subjected to flash column chromatography (40% ethyl acetate in hexane). The product was obtained as white solid with a yield of 510 mg, 60%. ^1^H NMR (400 MHz, Chloroform-*d*) δ 7.10 (d, *J* = 8.5 Hz, 2H), 6.83 (d, *J* = 8.6 Hz, 2H), 4.73 (s, 1H), 3.93 (t, *J* = 6.6 Hz, 2H), 3.81 (s, 1H), 3.60 (ddd, *J* = 46.0, 11.0, 4.6 Hz, 2H), 2.77 (d, *J* = 7.1 Hz, 2H), 2.27 (s, 1H), 1.89 – 1.68 (m, 2H), 1.55 – 1.26 (m, 13H), 0.93 (t, *J* = 7.0 Hz, 3H).

#### Synthesis of XML-10

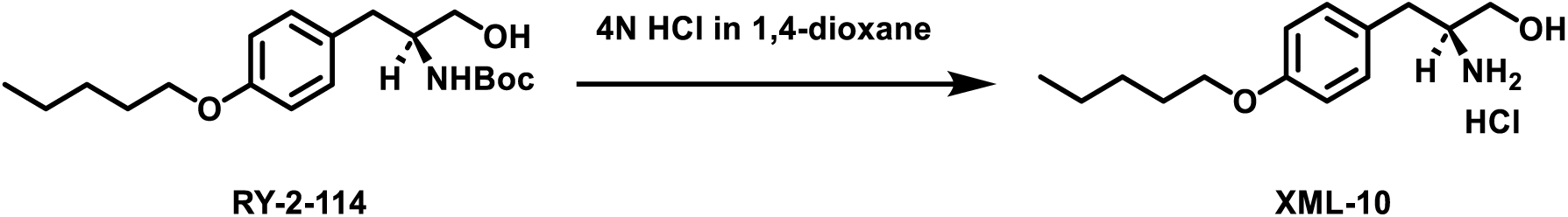

To a round-bottom flask was added RY-2-114 (400 mg, 1.2 mmol) and 4N HCl in 1,4-dioxane (2 mL). The reaction mixture was stirred for 2 h. Upon completion, the solvent was removed under reduced pressure. The residue was suspended in ethyl ether (5 mL) and filtered. The solid was washed with ethyl ether (10 mL) and dried under vacuum. The product was obtained as white solid with a yield of 260 mg, 80%. ^1^H NMR (400 MHz, Methanol-*d*4) δ 7.20 (d, *J* = 8.6 Hz, 2H), 6.91 (d, *J* = 8.6 Hz, 2H), 3.97 (t, *J* = 6.5 Hz, 2H), 3.72 (dd, *J* = 11.6, 3.6 Hz, 1H), 3.54 (dd, *J* = 11.6, 6.3 Hz, 1H), 3.47 – 3.37 (m, 1H), 2.90 (d, *J* = 7.4 Hz, 2H), 1.92 – 1.67 (m, 2H), 1.61 – 1.30 (m, 4H), 0.96 (t, *J* = 7.1 Hz, 3H). ^13^C NMR (101 MHz, Methanol-*d*4) δ 158.51, 129.96, 127.34, 114.58, 67.64, 60.00, 54.61, 34.21, 28.71, 27.99, 22.11, 12.97. HRMS-ESI: calculated for [C14H24NO2]^+^: 238.18016, found: 238.18009. UPLC: tR = 4.457 min, 83.45% (A1/B1); tR = 3.332 min, 86.54% (A2/B2).

#### Synthesis of RY-2-138

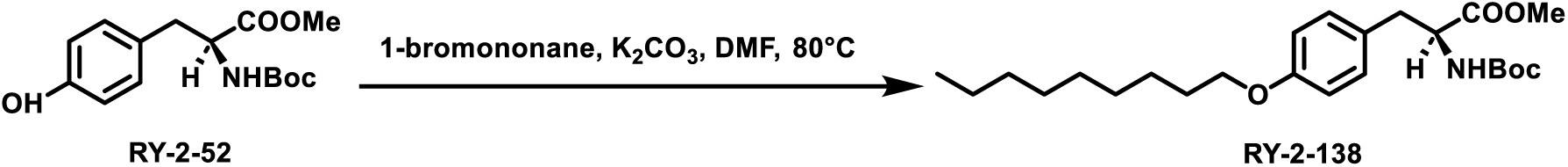

RY-2-52 (900 mg, 3 mmol, 1.0 eq) was dissolved in DMF (10 mL) followed by the addition of 1-bromononane (1242 mg, 6 mmol, 2.0 eq) and potassium carbonate (828 mg, 6 mmol, 2.0 eq). The reaction mixture was heated at 80°C and stirred overnight. Upon completion, the reaction mixture was quenched with water (50 mL) and extracted with ethyl acetate (50 mL x 3). The organic phase was washed with brine (50 mL x 3) and dried over sodium sulfate. The solvent was removed under reduced pressure and the residue was subjected to flash column chromatography (30% ethyl acetate in hexane). The product was obtained as white solid with a yield of 1.2 g, 93%. ^1^H NMR (400 MHz, Chloroform-*d*) δ 7.01 (d, *J* = 8.6 Hz, 2H), 6.81 (d, *J* = 8.6 Hz, 2H), 4.95 (d, *J* = 8.4 Hz, 1H), 4.53 (d, *J* = 7.0 Hz, 1H), 3.92 (t, *J* = 6.6 Hz, 2H), 3.71 (s, 3H), 3.21 – 2.90 (m, 2H), 1.92 – 1.65 (m, 2H), 1.52 – 1.13 (m, 22H), 1.02 – 0.74 (m, 3H).

#### Synthesis of RY-2-142

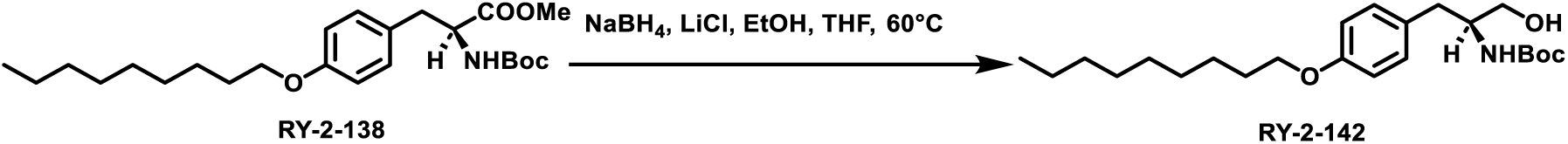

RY-2-138 (843 mg, 2.0 mmol, 1.0 eq) was dissolved in ethanol (10 mL) and THF (10 mL) followed by the addition of sodium borohydride (303 mg, 8 mmol, 4.0 eq) and lithium chloride (339 mg, 8 mmol, 4.0 eq). The reaction mixture was heated at 60°C and stirred overnight. Upon completion, the reaction mixture was slowly quenched with water (50 mL) and extracted with ethyl acetate (100 mL x 3). The organic phase was washed with brine (100 mL x 3) and dried over sodium sulfate. The solvent was removed under reduced pressure and the residue was subjected to flash column chromatography (40% ethyl acetate in hexane). The product was obtained as white solid with a yield of 680 mg, 86%. ^1^H NMR (400 MHz, Chloroform-*d*) δ 7.10 (d, *J* = 8.6 Hz, 2H), 6.83 (d, *J* = 8.6 Hz, 2H), 4.74 (s, 1H), 3.92 (t, *J* = 6.6 Hz, 2H), 3.81 (s, 1H), 3.72 – 3.46 (m, 2H), 2.77 (d, *J* = 7.2 Hz, 2H), 2.39 (s, 1H), 1.89 – 1.65 (m, 2H), 1.52 – 1.14 (m, 22H), 1.02 – 0.77 (m, 3H).

#### Synthesis of XML-11

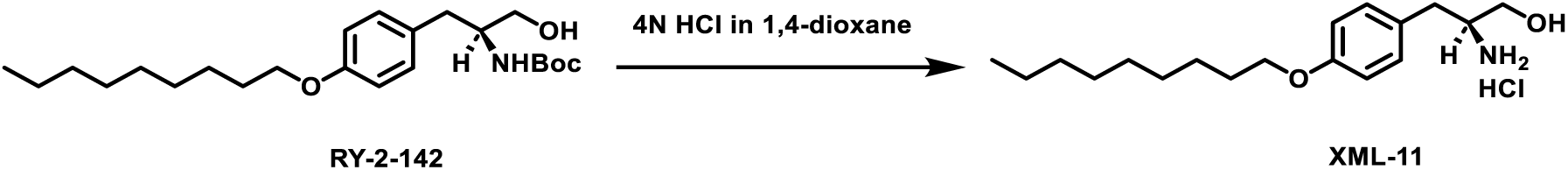

To a round-bottom flask was added RY-2-142 (400 mg, 1 mmol) and 4N HCl in 1,4-dioxane (2 mL). The reaction mixture was stirred for 2 h. Upon completion, the solvent was removed under reduced pressure. The residue was suspended in ethyl ether (5 mL) and filtered. The solid was washed with ethyl ether (10 mL) and dried under vacuum. The product was obtained as white solid with a yield of 283 mg, 84%. ^1^H NMR (400 MHz, Methanol-*d*4) δ 7.19 (d, *J* = 8.6 Hz, 2H), 6.90 (d, *J* = 8.6 Hz, 2H), 3.96 (t, *J* = 6.4 Hz, 2H), 3.71 (dd, *J* = 11.6, 3.6 Hz, 1H), 3.53 (dd, *J* = 11.6, 6.3 Hz, 1H), 3.47 – 3.34 (m, 1H), 2.89 (d, *J* = 7.4 Hz, 2H), 1.86 – 1.66 (m, 2H), 1.58 – 1.16 (m, 12H), 0.91 (t, *J* = 7.0 Hz, 3H). ^13^C NMR (101 MHz, Methanol-*d*4) δ 158.51, 129.96, 127.33, 114.58, 67.64, 60.01, 54.60, 34.22, 31.64, 29.29, 29.12, 29.00, 25.78, 22.33, 13.04. HRMS-ESI: calculated for [C18H32NO2]^+^: 294.24276, found: 294.24253. UPLC: tR = 5.429 min, >99% (A1/B1); tR = 4.165 min, 94.29% (A2/B2).

#### Synthesis of RY-2-140

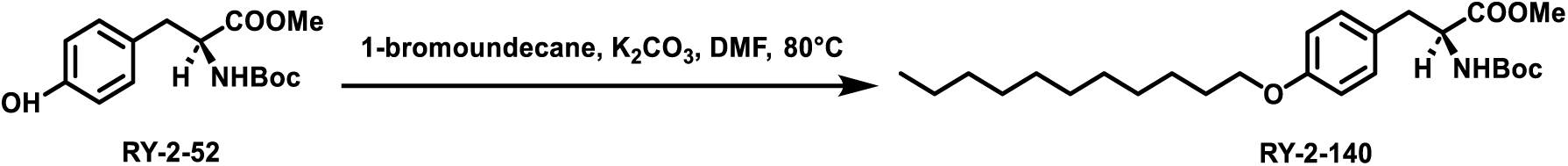

RY-2-52 (900 mg, 3 mmol, 1.0 eq) was dissolved in DMF (10 mL) followed by the addition of 1-bromoundecane (1411 mg, 6 mmol, 2.0 eq) and potassium carbonate (828 mg, 6 mmol, 2.0 eq). The reaction mixture was heated at 80°C and stirred overnight. Upon completion, the reaction mixture was quenched with water (50 mL) and extracted with ethyl acetate (50 mL x 3). The organic phase was washed with brine (50 mL x 3) and dried over sodium sulfate. The solvent was removed under reduced pressure and the residue was subjected to flash column chromatography (30% ethyl acetate in hexane). The product was obtained as white solid with a yield of 1.5 g, 96%. ^1^H NMR (400 MHz, Chloroform-*d*) δ 7.01 (d, *J* = 8.2 Hz, 2H), 6.81 (d, *J* = 8.6 Hz, 2H), 4.95 (d, *J* = 8.4 Hz, 1H), 4.53 (d, *J* = 7.2 Hz, 1H), 3.92 (t, *J* = 6.6 Hz, 2H), 3.71 (s, 3H), 3.13 – 2.92 (m, 2H), 1.85 – 1.70 (m, 2H), 1.51 – 1.06 (m, 35H), 0.88 (t, *J* = 6.6 Hz, 3H).

#### Synthesis of RY-2-143

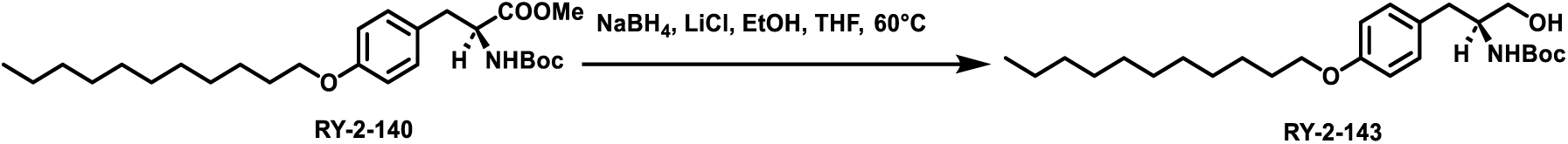

RY-2-140 (900 mg, 2.0 mmol, 1.0 eq) was dissolved in ethanol (10 mL) and THF (10 mL) followed by the addition of sodium borohydride (303 mg, 8 mmol, 4.0 eq) and lithium chloride (339 mg, 8 mmol, 4.0 eq). The reaction mixture was heated at 60°C and stirred overnight. Upon completion, the reaction mixture was slowly quenched with water (50 mL) and extracted with ethyl acetate (100 mL x 3). The organic phase was washed with brine (100 mL x 3) and dried over sodium sulfate. The solvent was removed under reduced pressure and the residue was subjected to flash column chromatography (60% ethyl acetate in hexane). The product was obtained as white solid with a yield of 600 mg, 71%. ^1^H NMR (400 MHz, Chloroform-*d*) δ 7.10 (d, *J* = 8.5 Hz, 2H), 6.83 (d, *J* = 8.5 Hz, 2H), 4.72 (s, 1H), 3.92 (t, *J* = 6.6 Hz, 2H), 3.81 (s, 1H), 3.60 (ddd, *J* = 46.4, 11.0, 4.6 Hz, 2H), 2.77 (d, *J* = 7.1 Hz, 2H), 2.21 (s, 1H), 1.87 – 1.67 (m, 2H), 1.54 – 1.11 (m, 26H), 0.88 (t, *J* = 6.7 Hz, 3H).

#### Synthesis of XML-12

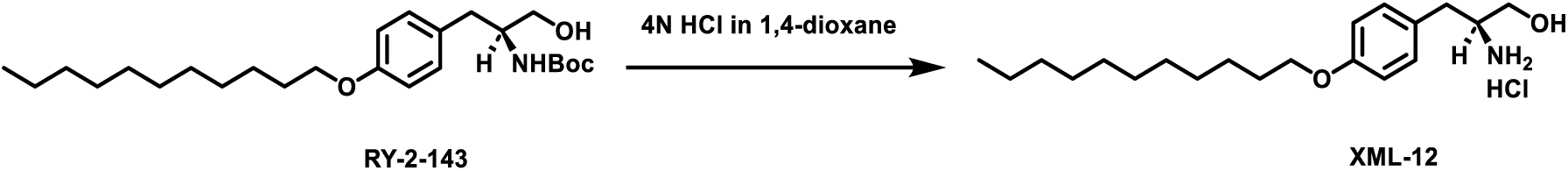

To a round-bottom flask was added RY-2-143 (350 mg, 0.8 mmol) and 4N HCl in 1,4-dioxane (2 mL). The reaction mixture was stirred for 2 h. Upon completion, the solvent was removed under reduced pressure. The residue was suspended in ethyl ether (5 mL) and filtered. The solid was washed with ethyl ether (10 mL) and dried under vacuum. The product was obtained as white solid with a yield of 220 mg, 74%. ^1^H NMR (400 MHz, Methanol-*d*4) δ 7.26 – 7.10 (m, 2H), 6.99 – 6.81 (m, 2H), 3.96 (t, *J* = 6.4 Hz, 2H), 3.71 (dd, *J* = 11.6, 3.6 Hz, 1H), 3.53 (dd, *J* = 11.6, 6.3 Hz, 1H), 3.47 – 3.36 (m, 1H), 2.89 (d, *J* = 7.4 Hz, 2H), 1.89 – 1.65 (m, 2H), 1.58 – 1.14 (m, 16H), 1.03 – 0.79 (m, 3H). ^13^C NMR (101 MHz, Methanol-*d*4) δ 158.51, 129.95, 127.31, 114.57, 67.62, 60.00, 54.59, 34.21, 31.67, 29.33, 29.11, 29.07, 29.01, 25.78, 22.34, 13.05. HRMS-ESI: calculated for [C20H36NO2]^+^: 322.27406, found: 322.27349. UPLC: tR = 5.718 min, >99% (A1/B1); tR = 4.542 min, >99% (A2/B2).

#### Synthesis of RY-2-137

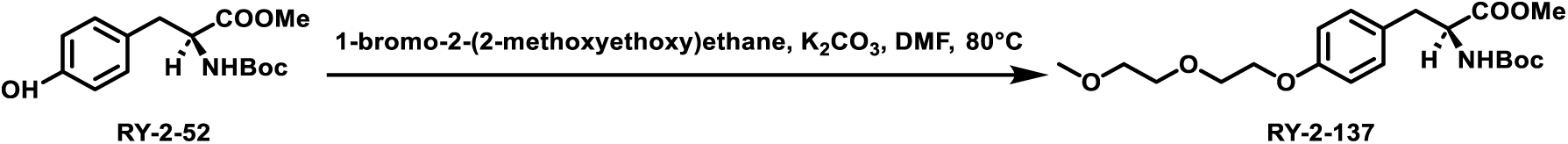

RY-2-52 (900 mg, 3 mmol, 1.0 eq) was dissolved in DMF (10 mL) followed by the addition of 1-bromo-2-(2-methoxyethoxy)ethane (1098 mg, 6 mmol, 2.0 eq) and potassium carbonate (828 mg, 6 mmol, 2.0 eq). The reaction mixture was heated at 80°C and stirred overnight. Upon completion, the reaction mixture was quenched with water (50 mL) and extracted with ethyl acetate (50 mL x 3). The organic phase was washed with brine (50 mL x 3) and dried over sodium sulfate. The solvent was removed under reduced pressure and the residue was subjected to flash column chromatography (40% ethyl acetate in hexane). The product was obtained as colorless oil with a yield of 980 mg, 82%. ^1^H NMR (400 MHz, Chloroform-*d*) δ 7.02 (d, *J* = 8.6 Hz, 2H), 6.84 (d, *J* = 8.6 Hz, 2H), 4.96 (d, *J* = 8.3 Hz, 1H), 4.53 (d, *J* = 7.0 Hz, 1H), 4.20 – 4.02 (m, 2H), 3.94 – 3.80 (m, 2H), 3.79 – 3.65 (m, 5H), 3.64 – 3.50 (m, 2H), 3.39 (s, 3H), 3.13 – 2.93 (m, 2H), 1.42 (s, 9H).

#### Synthesis of RY-2-141

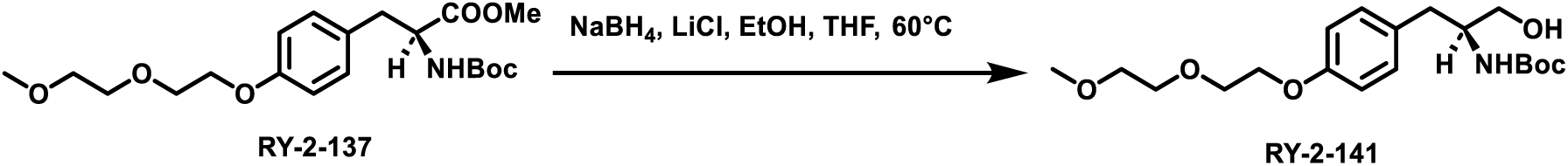

RY-2-137 (800 mg, 2.0 mmol, 1.0 eq) was dissolved in ethanol (10 mL) and THF (10 mL) followed by the addition of sodium borohydride (303 mg, 8 mmol, 4.0 eq) and lithium chloride (339 mg, 8 mmol, 4.0 eq). The reaction mixture was heated at 60°C and stirred overnight. Upon completion, the reaction mixture was slowly quenched with water (50 mL) and extracted with ethyl acetate (100 mL x 3). The organic phase was washed with brine (100 mL x 3) and dried over sodium sulfate. The solvent was removed under reduced pressure and the residue was subjected to flash column chromatography (60% ethyl acetate in hexane). The product was obtained as colorless oil with a yield of 670 mg, 91%. ^1^H NMR (400 MHz, Chloroform-*d*) δ 7.11 (d, *J* = 8.5 Hz, 2H), 6.84 (d, *J* = 8.6 Hz, 2H), 5.01 – 4.75 (m, 1H), 4.23 – 4.03 (m, 2H), 3.89 – 3.82 (m, 2H), 3.80 (s, 1H), 3.75 – 3.69 (m, 2H), 3.66 – 3.55 (m, 3H), 3.51 (dd, *J* = 11.0, 5.2 Hz, 1H), 3.46 (s, 3H), 3.39 (s, 3H), 2.76 (d, *J* = 7.1 Hz, 2H), 2.42 (s, 2H), 1.41 (s, 9H).

#### Synthesis of XML-13

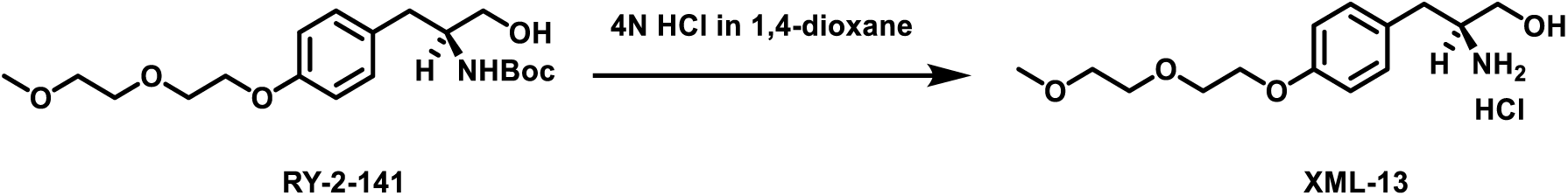

To a round-bottom flask was added RY-2-141 (400 mg, 1.1 mmol) and 4N HCl in 1,4-dioxane (3 mL). The reaction mixture was stirred for 2 h. Upon completion, the solvent was removed under reduced pressure. The residue was suspended in ethyl ether (5 mL) and filtered. The solid was washed with ethyl ether (10 mL) and dried under vacuum. The product was obtained as white solid with a yield of 220 mg, 66%. ^1^H NMR (400 MHz, Methanol-*d*4) δ 7.22 (d, *J* = 8.6 Hz, 2H), 6.95 (d, *J* = 8.6 Hz, 2H), 4.13 (t, *J* = 4.6 Hz, 2H), 3.84 (t, *J* = 4.7 Hz, 2H), 3.78 – 3.66 (m, 3H), 3.63 – 3.49 (m, 3H), 3.46 – 3.39 (m, 1H), 3.38 (s, 3H), 2.90 (d, *J* = 7.4 Hz, 2H). ^13^C NMR (101 MHz, Methanol-*d*4) δ 158.21, 130.01, 127.76, 114.71, 71.57, 70.14, 69.44, 67.25, 60.00, 57.71, 54.57, 34.20. HRMS-ESI: calculated for [C14H24NO4]^+^: 270.16998, found: 270.16959. UPLC: tR = 2.956 min, >99% (A1/B1); tR = 2.374 min, >99% (A2/B2).

#### Synthesis of RY-2-98

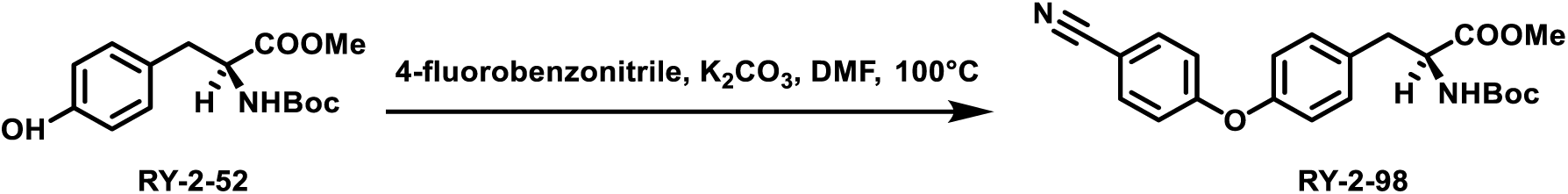

RY-2-52 (295 mg, 1 mmol, 1.0 eq) was dissolved in DMF (5 mL) followed by the addition of 4-fluorobenzonitrile (182 mg, 1.5 mmol, 1.5 eq) and potassium carbonate (276 mg, 1 mmol, 2.0 eq). The reaction mixture was heated at 100°C and stirred overnight. Upon completion, the reaction mixture was quenched with water (25 mL) and extracted with ethyl acetate (25 mL x 3). The organic phase was washed with brine (25 mL x 3) and dried over sodium sulfate. The solvent was removed under reduced pressure and the residue was subjected to flash column chromatography (35% ethyl acetate in hexane). The product was obtained as colorless oil with a yield of 96 mg, 24%. ^1^H NMR (400 MHz, Chloroform-*d*) δ 7.59 (d, *J* = 8.8 Hz, 2H), 7.18 (d, *J* = 8.5 Hz, 2H), 6.99 (d, *J* = 8.7 Hz, 4H), 5.06 (d, *J* = 8.3 Hz, 1H), 4.60 (d, *J* = 7.2 Hz, 1H), 3.74 (s, 3H), 3.09 (ddd, *J* = 55.5, 13.9, 6.1 Hz, 2H), 1.42 (s, 9H).

#### Synthesis of RY-2-106

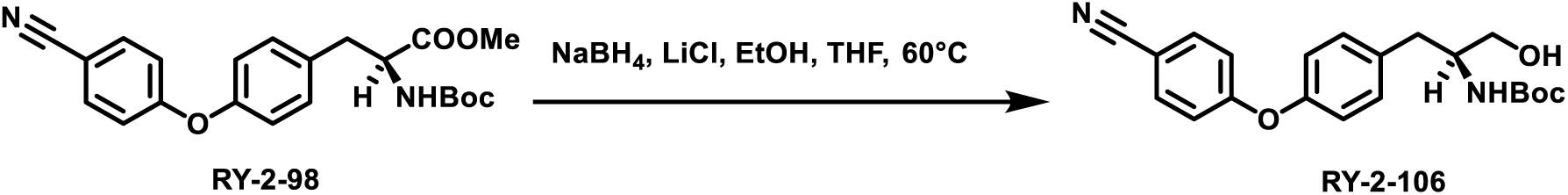

RY-2-98 (792 mg, 2.0 mmol, 1.0 eq) was dissolved in ethanol (10 mL) and THF (10 mL) followed by the addition of sodium borohydride (303 mg, 8 mmol, 4.0 eq) and lithium chloride (339 mg, 8 mmol, 4.0 eq). The reaction mixture was heated at 60°C and stirred overnight. Upon completion, the reaction mixture was slowly quenched with water (50 mL) and extracted with ethyl acetate (100 mL x 3). The organic phase was washed with brine (100 mL x 3) and dried over sodium sulfate. The solvent was removed under reduced pressure and the residue was subjected to flash column chromatography (70% ethyl acetate in hexane). The product was obtained as white solid with a yield of 535 mg, 73%. ^1^H NMR (400 MHz, Chloroform-*d*) δ 7.72 – 7.46 (m, 2H), 7.26 (t, *J* = 4.2 Hz, 2H), 7.12 – 6.88 (m, 4H), 4.78 (d, *J* = 8.2 Hz, 1H), 3.87 (s, 1H), 3.75 – 3.48 (m, 2H), 2.86 (d, *J* = 7.2 Hz, 2H), 2.08 (s, 1H), 1.42 (s, 9H).

#### Synthesis of XML-14

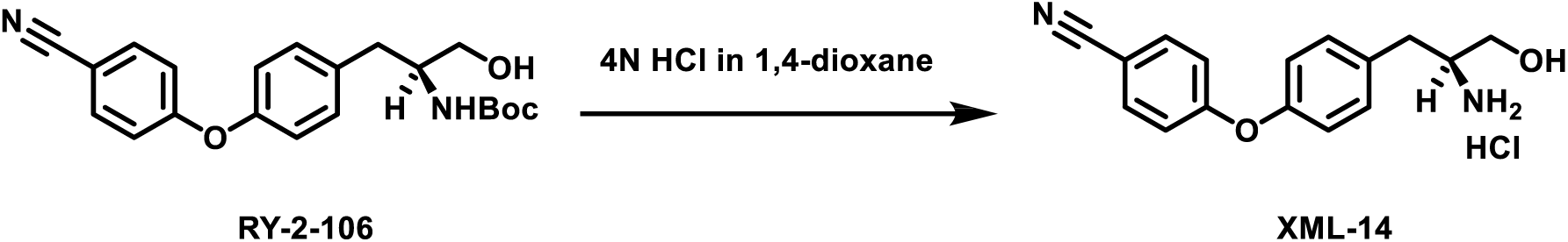

To a round-bottom flask was added RY-2-106 (200 mg, 0.5 mmol) and 4N HCl in 1,4-dioxane (5 mL). The reaction mixture was stirred for 2 h. Upon completion, the solvent was removed under reduced pressure. The residue was suspended in ethyl ether (5 mL) and filtered. The solid was washed with ethyl ether (10 mL) and dried under vacuum. The product was obtained as white solid with a yield of 110 mg, 66%. ^1^H NMR (400 MHz, Methanol-*d*4) δ 7.71 (d, *J* = 8.9 Hz, 2H), 7.42 (d, *J* = 8.5 Hz, 2H), 7.10 (dd, *J* = 8.7, 6.7 Hz, 4H), 3.77 (dd, *J* = 11.5, 3.6 Hz, 1H), 3.60 (dd, *J* = 11.5, 6.1 Hz, 1H), 3.56 – 3.47 (m, 1H), 3.02 (d, *J* = 7.4 Hz, 2H). ^13^C NMR (101 MHz, Methanol-*d*4) δ 161.76, 154.20, 134.09, 132.77, 131.00, 120.53, 118.29, 117.78, 59.95, 54.39, 34.32. HRMS-ESI: calculated for [C16H17N2O2]^+^: 269.12845, found: 269.12804. UPLC: tR = 3.790 min, >99% (A1/B1); tR = 2.957 min, >99% (A2/B2).

#### Synthesis of RY-2-135

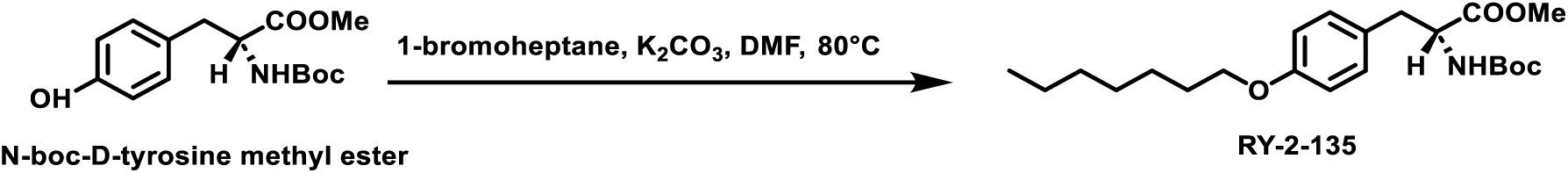

N-boc-D-tyrosine methyl ester (2953 mg, 10 mmol, 1.0 eq) was dissolved in DMF (50 mL) followed by the addition of 1-bromoheptane (3582 mg, 20 mmol, 2.0 eq) and potassium carbonate (2760 mg, 20 mmol, 2.0 eq). The reaction mixture was heated at 80°C and stirred overnight. Upon completion, the reaction mixture was quenched with water (150 mL) and extracted with ethyl acetate (150 mL x 3). The organic phase was washed with brine (150 mL x 3) and dried over sodium sulfate. The solvent was removed under reduced pressure and the residue was subjected to flash column chromatography (30% ethyl acetate in hexane). The product was obtained as white solid with a yield of 3.5 g, 89%. ^1^H NMR (400 MHz, Chloroform-*d*) δ 7.01 (d, *J* = 8.4 Hz, 2H), 6.81 (d, *J* = 8.6 Hz, 2H), 4.95 (d, *J* = 8.4 Hz, 1H), 4.53 (d, *J* = 7.1 Hz, 1H), 3.92 (t, *J* = 6.6 Hz, 2H), 3.71 (s, 3H), 3.13 – 2.93 (m, 2H), 1.88 – 1.69 (m, 2H), 1.53 – 1.14 (m, 19H), 1.01 – 0.76 (m, 3H).

#### Synthesis of RY-2-144

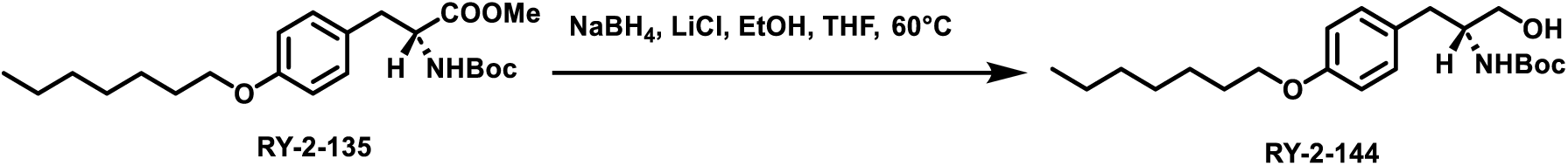

RY-2-135 (800 mg, 2 mmol, 1.0 eq) was dissolved in ethanol (10 mL) and THF (10 mL) followed by the addition of sodium borohydride (303 mg, 8 mmol, 4.0 eq) and lithium chloride (339 mg, 8 mmol, 4.0 eq). The reaction mixture was heated at 60°C and stirred overnight. Upon completion, the reaction mixture was slowly quenched with water (50 mL) and extracted with ethyl acetate (100 mL x 3). The organic phase was washed with brine (100 mL x 3) and dried over sodium sulfate. The solvent was removed under reduced pressure and the residue was subjected to flash column chromatography (40% ethyl acetate in hexane). The product was obtained as white solid with a yield of 680 mg, 93%. ^1^H NMR (400 MHz, Chloroform-*d*) δ 7.10 (d, *J* = 8.5 Hz, 2H), 6.83 (d, *J* = 8.6 Hz, 2H), 4.69 (s, 1H), 3.93 (t, *J* = 6.6 Hz, 2H), 3.81 (s, 1H), 3.60 (ddd, *J* = 47.1, 11.0, 4.5 Hz, 2H), 2.77 (d, *J* = 7.1 Hz, 2H), 1.94 (s, 1H), 1.83 – 1.70 (m, 2H), 1.54 – 1.20 (m, 17H), 1.00 – 0.81 (m, 3H).

#### Synthesis of XML-3D

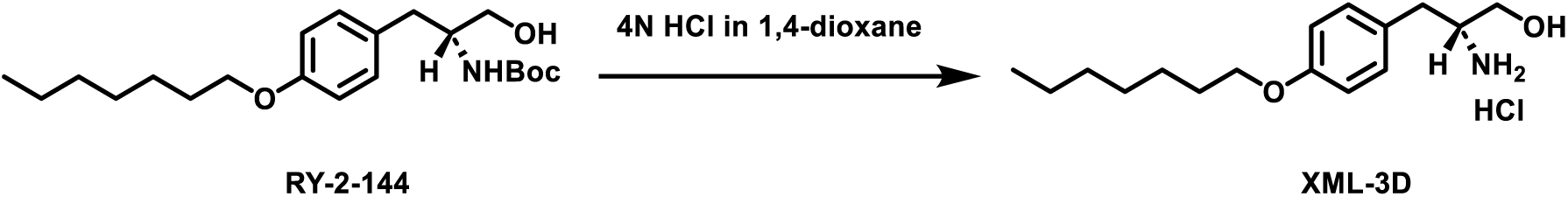

To a round-bottom flask was added RY-2-144 (400 mg, 1.1 mmol) and 4N HCl in 1,4-dioxane (3 mL). The reaction mixture was stirred for 2 h. Upon completion, the solvent was removed under reduced pressure. The residue was suspended in ethyl ether (5 mL) and filtered. The solid was washed with ethyl ether (10 mL) and dried under vacuum. The product was obtained as white solid with a yield of 230 mg, 70%. ^1^H NMR (400 MHz, Methanol-*d*4) δ 7.21 (d, *J* = 8.6 Hz, 2H), 6.92 (d, *J* = 8.6 Hz, 2H), 3.98 (t, *J* = 6.4 Hz, 2H), 3.73 (dd, *J* = 11.6, 3.5 Hz, 1H), 3.55 (dd, *J* = 11.6, 6.3 Hz, 1H), 3.50 – 3.37 (m, 1H), 2.91 (d, *J* = 7.4 Hz, 2H), 1.91 – 1.70 (m, 2H), 1.60 – 1.25 (m, 8H), 0.94 (t, *J* = 6.7 Hz, 2H). ^13^C NMR (101 MHz, Methanol-*d*4) δ 158.51, 129.96, 127.34, 114.58, 67.65, 60.00, 54.61, 34.21, 31.59, 29.02, 28.80, 25.75, 22.27, 13.02. HRMS-ESI: calculated for [C16H28NO2]^+^: 266.21146, found: 266.21118. UPLC: tR = 5.030 min, >99% (A1/B1); tR = 3.773 min, >99% (A2/B2).

### 2. Cell Lines and Cell Culture

DLBCL cell lines (Riva and Pfeiffer), MCL cell line (Granta-519, Jeko-1, Mino-1, UPN1), and human leukemia cell lines (MV4-11, HL-60, MEC-1) were cultured in standard RPMI-1640 medium (catalog number: 21870084, Thermo Fisher Scientific) supplemented with 10% Fetal bovine serum (FBS, catalog number: F4135, Sigma-Aldrich). All cells were maintained in a humidified incubator at 37 °C with 5% CO2.

### 3. Viability Assays

100,000 cells in complete cell culture medium (RPMI-1640 + 10% FBS) were seeded in 96-well plates (75 μL per well). Compounds were dissolved in DMSO and diluted (250X) in complete cell culture medium. 25 μL of each treatment-containing medium were added to each well (4X dilution). The complete cell culture medium with DMSO treatment was added in vehicle wells, and the complete cell culture medium without treatment was added in control wells. Wells containing the complete cell culture medium without cells served as background. The outer wells of the 96-well plates were filled with 200 μL of the complete cell culture medium to reduce evaporation from inner wells. Each compound was tested at eight different concentrations with three replicates. The treated cells were incubated 72 h at 37°C and 5% CO2. Cell viability was measured after 72 h incubation by the MTS assay (24h incubation for Jeko-1, Mino-1 and UPN1). 20 μL of the MTS reagent (Catalog number: G3580, Promega) was added to each well and the cells were incubated at the same condition for 4 h. The absorbance was measured at 490 nm using BioTek Synergy HTX Multimode Reader. The average absorbance value of background wells was subtracted from absorbance value of each vehicle and treatment well. The percentage cell viability in each treatment well was determined by normalizing the absorbance of each treatment well to the average absorbance value of vehicle wells. The data fitted in non-linear regression curves using GraphPad Prism 9 (GraphPad Software, La Jolla, CA, USA).

### 4. PP2A Activity Assay

#### Lysate preparation

MV4-11 cells were treated with three different XML-3D or OSU-2S concentrations (1.5–5 μM) for 4 hours with three technical replicates. For inhibitor studies, Jeko-1 cells were pretreated with 5 nM okadaic acid for 2h prior to treatment with 5 µM XML-3D or DMSO vehicle for 6h with three technical replicates. Phosphatase activity was measured using the PP2A immunoprecipitation phosphatase assay kit from MilliporeSigma (catalog number: 17-313). Cells were washed in phosphate-free buffer to remove any free phosphate (10 mM HEPES buffer, 100 mM NaCl), then were suspended in 100 µL phosphate-free and low-detergent lysis buffer (10 mM HEPES, 1% Nonidet P-40, 0.5 M NaCl, 10% Glycerol, 1 mM PMSF and 1 mM sigma), incubated for 10 mins at 4°C then sonicated (15 secs on, 15 secs off three times).

#### PP2A Immunoprecipitation and activity assay

100 µg lysates were immunoprecipitated with either mouse anti-PP2Ac or corresponding mouse IgG2bƙ isotype control mixed with 40 µg protein A agarose beads diluted to 500 µL with pNPP Ser/Thr assay buffer (50 mM Tris-HCl, pH 7.0 and 100 µM CaCl2) for 2h at 4°C on a rotator. Beads with the immunocomplex were subsequently washed three times with TBS then with pNPP Ser/Thr assay buffer. Half of the beads were used in the activity assay, while 2x protein loading dye was added to the other half to run a Western blot and quantify PP2Ac protein expression for normalization of PP2A activity. Beads used in the activity assay were mixed with phosphopeptide (750 µM), diluted to 80 µL with pNPP ser/thr assay buffer and the reaction was incubated for 10 minutes at 30°C in a shaking incubator. Beads were centrifuged briefly and the supernatant was diluted 5X with 100 µL malachite green and incubated for 10 mins at room temperature before measuring the absorbance at 650 nm using BioTek Synergy HTX Multimode Reader. A free phosphate standard curve was used to calculate the free phosphate concentration in the samples.

#### Western blot for PP2Ac quantification

Proteins were resolved in an 8% tris-glycine gel, transferred to a nitrocellulose membrane (catalog number: 1704159, Bio-Rad), then blocked with 5% non-fat dry milk in Trisbuffered saline-Tween-20 (TBST) for 1h at room temperature. Primary rabbit monoclonal anti-PP2Ac (1:1000, catalogue number: 2259, Cell Signaling Technology, overnight at 4°C) and secondary goat anti-rabbit IgG (H+L) HRP conjugated (1:3000, catalogue number: 170-6515, Bio-Rad, 1h room temperature) in 5% non-fat dry milk in TBST were used to stain the blots overnight at 4°C and 1h at room temperature respectively, which were visualized using a ChemiDoc Imaging System (Bio-Rad) and quantified using the image J software.

### 5. Co-Immunoprecipitation

### Antibody/beads conjugation/crosslinking

Rabbit monoclonal anti-PP2Ac antibody (catalog number: 2259, 1:25 dilution) or corresponding isotype control (rabbit monoclonal IgG, catalogue number: AB172730, Abcam) and 40 µg protein A agarose beads (catalogue number: 16-125 MilliporeSigma) were added to 500 µL immunoprecipitation (IP) lysis buffer (20mM Tris, 500mM NaCl, 1% Nonidet P-40, 1mM EDTA, pH 7.5). The reaction was incubated at 4°C for 4h on a rotator to form the IgG/beads complex, then washed twice with conjugation buffer (20 mM Sodium Phosphate, 0.15M NaCl (pH 7-9)). Subsequently, the IgG/beads complex was crosslinked in 250 µL 5 mM BS^3^ irreversible crosslinker solution (catalogue number: 21580, Thermo Fisher Scientific) for 30 mins, then quenched with 12.5 µL quenching buffer (1M Tris HCl (pH 7.5)) for 15 mins at room temperature on a rotator. The crosslinked complex was washed three times with PBST.

### Immunoprecipitation

Jeko-1 cells and MV4-11 cells were treated with 1 µM XML-3D or vehicle for 6h and 2 µM XML-3D or vehicle for 4h respectively. Lysates were prepared by incubating cells (60e6) in IP lysis buffer for 10 mins at 4°C. 3 mg lysate was precleared twice with a crosslinked rabbit IgG control/beads complex for 1h on a rotator at 4°C twice. Half of the precleared lysate was incubated with the anti-PP2Ac/beads complex, and the other half was incubated with the rabbit IgG control/beads complex overnight at 4°C on a rotator. The immunoprecipitated samples were washed three times with IP lysis buffer before adding 30 µL of 2x protein loading dye. Western blot was performed as described above. Primary rabbit monoclonal anti-PP2Ac (1:1000, catalogue number: 2259, overnight at 4°C) and rabbit monoclonal anti-SET (1:1000, catalog number: MA5-34662, Fisher Scientific, overnight at 4°C), and secondary antibody Veriblot HRP conjugated (1:250, catalog number AB131366, Abcam, 2h at room temperature) that only recognizes antibodies in their native conformation were used to stain the blots. All primary and secondary antibodies were dissolved in 5% non-fat dry milk in TBST. PP2Ac and SET protein levels were also quantified in the input.

## Reference

(1) Deribe, Y. L.; Pawson, T.; Dikic, I. Post-translational modifications in signal integration. Nat. Struct. Mol. Biol. 2010, 17, 666–672. DOI: 10.1038/nsmb.1842.

(2) Ardito, F.; Giuliani, M.; Perrone, D.; Troiano, G.; Lo Muzio, L. The crucial role of protein phosphorylation in cell signaling and its use as targeted therapy (Review). Int. J. Mol. Med. 2017, 40, 271–280. DOI: 10.3892/ijmm.2017.3036.

(3) Tonks, N. K. Protein tyrosine phosphatases – from housekeeping enzymes to master regulators of signal transduction. FEBS J. 2013, 280, 346–378. DOI: 10.1111/febs.12077.

(4) Roskoski, R. A historical overview of protein kinases and their targeted small molecule inhibitors. Pharmacol. Res. 2015, 100, 1–23. DOI: 10.1016/j.phrs.2015.07.010.

(5) Roskoski, R. Properties of FDA-approved small molecule protein kinase inhibitors: A 2024 update. Pharmacol. Res. 2024, 200, 107059. DOI: 10.1016/j.phrs.2024.107059.

(6) O’Connor, C. M.; Perl, A.; Leonard, D.; Sangodkar, J.; Narla, G. Therapeutic targeting of PP2A. Int. J. Biochem. Cell Biol. 2018, 96, 182–193. DOI: 10.1016/j.biocel.2017.10.008.

(7) Brautigan, D. L.; Shenolikar, S. Protein Serine/Threonine Phosphatases: Keys to Unlocking Regulators and Substrates. Annu. Rev. Biochem. 2018, 87, 921–964. DOI: 10.1146/annurev-biochem-062917-012448.

(8) Stanford, S. M.; Bottini, N. Targeting protein phosphatases in cancer immunotherapy and autoimmune disorders. Nat. Rev. Drug Discovery 2023, 22, 273–294. DOI: 10.1038/s41573-022-00618-w.

(9) Shi, Y. Serine/Threonine Phosphatases: Mechanism through Structure. Cell 2009, 139, 468–484. DOI: 10.1016/j.cell.2009.10.006.

(10) Seshacharyulu, P.; Pandey, P.; Datta, K.; Batra, S. K. Phosphatase: PP2A structural importance, regulation and its aberrant expression in cancer. Cancer Lett. 2013, 335, 9–18. DOI: 10.1016/j.canlet.2013.02.036.

(11) Zolnierowicz, S.; Csortos, C.; Bondor, J.; Verin, A.; Mumby, M. C.; DePaoli-Roach, A. A. Diversity in the Regulatory B-Subunits of Protein Phosphatase 2A: Identification of a Novel Isoform Highly Expressed in Brain. Biochemistry 1994, 33, 11858–11867. DOI: 10.1021/bi00205a023.

(12) Hemmings, B. A.; Adams-Pearson, C.; Maurer, F.; Mueller, P.; Goris, J.; Merlevede, W.; Hofsteenge, J.; Stone, S. R. α- and β-forms of the 65-kDa subunit of protein phosphatase 2A have a similar 39 amino acid repeating structure. Biochemistry 1990, 29, 3166–3173. DOI: 10.1021/bi00465a002.

(13) Yang, G.; Murashige, D. S.; Humphrey, S. J.; James, D. E. A Positive Feedback Loop between Akt and mTORC2 via SIN1 Phosphorylation. Cell Rep. 2015, 12, 937–943. DOI: 10.1016/j.celrep.2015.06.070.

(14) Manning, B. D.; Toker, A. AKT/PKB Signaling: Navigating the Network. Cell 2017, 169, 381–405. DOI: 10.1016/j.cell.2017.03.034.

(15) Kuo, Y.-C.; Huang, K.-Y.; Yang, C.-H.; Yang, Y.-S.; Lee, W.-Y.; Chiang, C.-W. Regulation of Phosphorylation of Thr-308 of Akt, Cell Proliferation, and Survival by the B55α Regulatory Subunit Targeting of the Protein Phosphatase 2A Holoenzyme to Akt. J. Biol. Chem. 2008, 283, 1882–1892. DOI: 10.1074/jbc.M709585200.

(16) Liao, Y.; Hung, M.-C. Physiological regulation of Akt activity and stability. Am. J. Transl. Res. 2010, 2, 19–42.

(17) Migliaccio, N.; Sanges, C.; Ruggiero, I.; Martucci, N. M.; Rippa, E.; Arcari, P.; Lamberti, A. Raf kinases in signal transduction and interaction with translation machinery. Biomol. Concepts 2013, 4, 391–399. DOI: 10.1515/bmc-2013-0007.

(18) Zang, M.; Gong, J.; Luo, L.; Zhou, J.; Xiang, X.; Huang, W.; Huang, Q.; Luo, X.; Olbrot, M.; Peng, Y.; et al. Characterization of Ser338 Phosphorylation for Raf-1 Activation. J. Biol. Chem. 2008, 283, 31429–31437. DOI: 10.1074/jbc.M801066200.

(19) Janssens, V.; Goris, J.; Van Hoof, C. PP2A: the expected tumor suppressor. Curr. Opin. Genet. Dev. 2005, 15, 34–41. DOI: 10.1016/j.gde.2004.12.004.

(20) Cristóbal, I.; Rincón, R.; Manso, R.; Caramés, C.; Zazo, S.; Madoz-Gúrpide, J.; Rojo, F.; García-Foncillas, J. Deregulation of the PP2A Inhibitor SET Shows Promising Therapeutic Implications and Determines Poor Clinical Outcome in Patients with Metastatic Colorectal Cancer. Clin. Cancer Res. 2015, 21, 347–356. DOI: 10.1158/1078-0432.CCR-14-0801.

(21) Calin, G. A.; di Iasio, M. G.; Caprini, E.; Vorechovsky, I.; Natali, P. G.; Sozzi, G.; Croce, C. M.; Barbanti-Brodano, G.; Russo, G.; Negrini, M. Low frequency of alterations of the α (PPP2R1A) and β (PPP2R1B) isoforms of the subunit A of the serine-threonine phosphatase 2A in human neoplasms. Oncogene 2000, 19, 1191–1195. DOI: 10.1038/sj.onc.1203388.

(22) Colella, S.; Ohgaki, H.; Ruediger, R.; Yang, F.; Nakamura, M.; Fujisawa, H.; Kleihues, P.; Walter, G. Reduced expression of the Aα subunit of protein phosphatase 2A in human gliomas in the absence of mutations in the Aα and Aβ subunit genes. Int. J. Cancer 2001, 93, 798–804. DOI: 10.1002/ijc.1404.

(23) Li, M.; Guo, H.; Damuni, Z. Purification and Characterization of Two Potent Heat-Stable Protein Inhibitors of Protein Phosphatase 2A from Bovine Kidney. Biochemistry 1995, 34, 1988–1996. DOI: 10.1021/bi00006a020.

(24) Khanna, A.; Pimanda, J. E.; Westermarck, J. Cancerous Inhibitor of Protein Phosphatase 2A, an Emerging Human Oncoprotein and a Potential Cancer Therapy Target. Cancer Res. 2013, 73, 6548–6553. DOI: 10.1158/0008-5472.CAN-13-1088.

(25) Cristóbal, I.; Garcia-Ortiz, L.; Calleros, L.; et al. Overexpression of SET is a recurrent event associated with poor outcome and contributes to protein phosphatase 2A inhibition in acute myeloid leukemia. Haematologica 2012, 97, 543–550. DOI: 10.3324/haematol.2011.050542.

(26) Zonta, F.; Pagano, M. A.; Trentin, L.; Tibaldi, E.; Frezzato, F.; Trimarco, V.; Facco, M.; Zagotto, G.; Pavan, V.; Ribaudo, G.; Bordin, L.; Semenzato, G. C.; Brunati, A. M. Lyn Sustains Oncogenic Signaling in Chronic Lymphocytic Leukemia by Strengthening SET-Mediated Inhibition of PP2A. Blood 2015, 125, 3747–3755. DOI: 10.1182/blood-2014-12-619155.

(27) Liu, H.; Gu, Y.; Wang, H.; Yin, J.; Zheng, G.; Zhang, Z.; Lu, M.; Wang, C.; He, Z. Overexpression of PP2A inhibitor SET oncoprotein is associated with tumor progression and poor prognosis in human non-small cell lung cancer. Oncotarget 2015, 6, 14913–14925. DOI: 10.18632/oncotarget.3818.

(28) Cristóbal, I.; Torrejón, B.; Rubio, J.; Santos, A.; Pedregal, M.; Caramés, C.; Zazo, S.; Luque, M.; Sanz-Alvarez, M.; Madoz-Gúrpide, J.; et al. Deregulation of SET is Associated with Tumor Progression and Predicts Adverse Outcome in Patients with Early-Stage Colorectal Cancer. J. Clin. Med. 2019, 8, 346. DOI: 10.3390/jcm8030346.

(29) Hung, M. H.; Chen, Y. L.; Chu, P. Y.; Shih, C. T.; Yu, H. C.; Tai, W. T.; Shiau, C. W.; Chen, K. F. Upregulation of the oncoprotein SET determines poor clinical outcomes in hepatocellular carcinoma and shows therapeutic potential. Oncogene 2016, 35, 4891–4902. DOI: 10.1038/onc.2015.455.

(30) Hu, X.; Garcia, C.; Fazli, L.; Gleave, M.; Vitek, M. P.; Jansen, M.; Christensen, D.; Mulholland, D. J. Inhibition of Pten deficient Castration Resistant Prostate Cancer by Targeting of the SET-PP2A Signaling axis. Sci. Rep. 2015, 5, 15182. DOI: 10.1038/srep15182.

(31) Liu, C.-Y.; Huang, T.-T.; Chen, Y.-T.; Chen, J.-L.; Chu, P.-Y.; Huang, C.-T.; Wang, W.-L.; Lau, K.-Y.; Dai, M.-S.; Shiau, C.-W.; et al. Targeting SET to restore PP2A activity disrupts an oncogenic CIP2A-feedforward loop and impairs triple negative breast cancer progression. EBioMedicine 2019, 40, 263–275. DOI: 10.1016/j.ebiom.2019.01.023.

(32) Chalfant, C. E.; Kishikawa, K.; Mumby, M. C.; Kamibayashi, C.; Bielawska, A.; Hannun, Y. A. Long Chain Ceramides Activate Protein Phosphatase-1 and Protein Phosphatase-2A: Activation Is Stereospecific and Regulated by Phosphatidic Acid. J. Biol. Chem. 1999, 274, 20313–20317. DOI: 10.1074/jbc.274.29.20313.

(33) Dobrowsky, R. T.; Kamibayashi, C.; Mumby, M. C.; Hannun, Y. A. Ceramide activates heterotrimeric protein phosphatase 2A. J. Biol. Chem. 1993, 268, 15523–15530.

(34) Mukhopadhyay, A.; Saddoughi, S. A.; Song, P.; Sultan, I.; Ponnusamy, S.; Senkal, C. E.; Snook, C. F.; Arnold, H. K.; Sears, R. C.; Hannun, Y. A.; et al. Direct interaction between the inhibitor 2 and ceramide via sphingolipid-protein binding is involved in the regulation of protein phosphatase 2A activity and signaling. FASEB J. 2009, 23, 751–763. DOI: 10.1096/fj.08-120550.

(35) Brinkmann, V. FTY720 (fingolimod) in Multiple Sclerosis: therapeutic effects in the immune and the central nervous system. Br. J. Pharmacol. 2009, 158, 1173–1182. DOI: 10.1111/j.1476-5381.2009.00451.x.

(36) Chen, L.; Luo, L.-F.; Lu, J.; Li, L.; Liu, Y.-F.; Wang, J.; Liu, H.; Song, H.; Jiang, H.; Chen, S.-J.; et al. FTY720 Induces Apoptosis of M2 Subtype Acute Myeloid Leukemia Cells by Targeting Sphingolipid Metabolism and Increasing Endogenous Ceramide Levels. PLoS One 2014, 9, e103033. DOI: 10.1371/journal.pone.0103033.

(37) Liang, J.; Nagahashi, M.; Kim, E. Y.; Harikumar, K. B.; Yamada, A.; Huang, W.-C.; Hait, N. C.; Allegood, J. C.; Price, M. M.; Avni, D.; et al. Sphingosine-1-Phosphate Links Persistent STAT3 Activation, Chronic Intestinal Inflammation, and Development of Colitis-Associated Cancer. Cancer Cell 2013, 23, 107–120. DOI: 10.1016/j.ccr.2012.11.013.

(38) Matsuoka, Y.; Nagahara, Y.; Ikekita, M.; Shinomiya, T. A novel immunosuppressive agent FTY720 induced Akt dephosphorylation in leukemia cells. Br. J. Pharmacol. 2003, 138, 1303–1312. DOI: 10.1038/sj.bjp.0705181.

(39) Neviani, P.; Santhanam, R.; Oaks, J. J.; Eiring, A. M.; Notari, M.; Blaser, B. W.; Liu, S.; Trotta, R.; Muthusamy, N.; Gambacorti-Passerini, C.; et al. FTY720, a new alternative for treating blast crisis chronic myelogenous leukemia and Philadelphia chromosome-positive acute lymphocytic leukemia. J. Clin. Invest. 2007, 117, 2408–2421. DOI: 10.1172/JCI31095.

(40) Saddoughi, S. A.; Gencer, S.; Peterson, Y. K.; Ward, K. E.; Mukhopadhyay, A.; Oaks, J.; Bielawski, J.; Szulc, Z. M.; Thomas, R. J.; Selvam, S. P.; et al. Sphingosine analogue drug FTY720 targets I2PP2A/SET and mediates lung tumour suppression via activation of PP2A-RIPK1-dependent necroptosis. EMBO Mol. Med. 2013, 5, 105–121. DOI: 10.1002/emmm.201201283.

(41) Shouse, G.; de Necochea-Campion, R.; Mirshahidi, S.; Liu, X.; Chen, C.-S. Novel B55α-PP2A mutations in AML promote AKT T308 phosphorylation and sensitivity to AKT inhibitor-induced growth arrest. Oncotarget 2016, 7, 61081–61092. DOI: 10.18632/oncotarget.11268.

(42) Kiuchi, M.; Adachi, K.; Kohara, T.; Teshima, K.; Masubuchi, Y.; Mishina, T.; Fujita, T. Synthesis and Biological Evaluation of 2,2-Disubstituted 2-Aminoethanols: Analogues of FTY720. Bioorg. Med. Chem. Lett. 1998, 8, 101–106. DOI: 10.1016/S0960-894X(97)10116-2.

(43) Mani, R.; Mao, Y.; Frissora, F. W.; et al. ROR1-targeted delivery of OSU-2S exerts cytotoxicity in mantle cell lymphoma. Exp. Hematol. 2015, 43, 770–774.e2. DOI: 10.1016/j.exphem.2015.04.008.

(44) Camp, S. M.; Chiang, E. T.; Moreland, J. G.; et al. Pulmonary Endothelial Cell Barrier Enhancement by Novel FTY720 Analogs. Chem. Phys. Lipids 2016, 194, 85–93. DOI: 10.1016/j.chemphyslip.2015.10.004.

(45) Toop, H. D.; Dun, M. D.; Ross, B. K.; Flanagan, H. M.; Verrills, N. M.; Morris, J. C. Development of Novel PP2A Activators for Use in AML. Org. Biomol. Chem. 2016, 14, 4605–4616. DOI: 10.1039/C6OB00556J.

(46) Vicente, C.; Arriazu, E.; Martínez-Balsalobre, E.; Peris, I.; Marcotegui, N.; García-Ramírez, P.; Pippa, R.; Rabal, O.; Oyarzabal, J.; Guruceaga, E.; Prósper, F.; Mateos, M. C.; Cayuela, M. L.; Odero, M. D. A Novel FTY720 Analogue Targets SET-PP2A Interaction and Inhibits Growth of Acute Myeloid Leukemia Cells without Inducing Cardiac Toxicity. ACS Med. Chem. Lett. 2020, 11, 468–480. DOI: 10.1021/acsmedchemlett.0c00077.

(47) Johnson, H.; et al. Altering phosphorylation in cancer through PP2A modifiers. Cancer Cell Int. 2024, 24, 11. DOI: 10.1186/s12935-023-03193-1.

